# A spatially resolved multiomic single-cell atlas of soybean development

**DOI:** 10.1101/2024.07.03.601616

**Authors:** Xuan Zhang, Ziliang Luo, Alexandre P. Marand, Haidong Yan, Hosung Jang, Sohyun Bang, John P. Mendieta, Mark A.A. Minow, Robert J. Schmitz

**Affiliations:** Department of Genetics, University of Georgia, Athens, GA, USA; Department of Molecular, Cellular, and Development Biology, University of Michigan, Ann Arbor, MI, USA; Institute of Bioinformatics, University of Georgia, Athens, GA, USA; College of Grassland Science and Technology, Sichuan Agricultural University, Chengdu, China

## Abstract

*Cis*-regulatory elements (CREs) precisely control spatiotemporal gene expression in cells. Using a spatially resolved single-cell atlas of gene expression with chromatin accessibility across ten soybean tissues, we identified 103 distinct cell types and 303,199 accessible chromatin regions (ACRs). Nearly 40% of the ACRs showed cell-type-specific patterns and were enriched for transcription factor (TF) motifs defining diverse cell identities. We identified *de novo* enriched TF motifs and explored conservation of gene regulatory networks underpinning legume symbiotic nitrogen fixation. With comprehensive developmental trajectories for endosperm and embryo, we uncovered the functional transition of the three sub-cell types of endosperm, identified 13 sucrose transporters sharing the DOF11 motif that were co-up-regulated in late peripheral endosperm and identified key embryo cell-type specification regulators during embryogenesis, including a homeobox TF that promotes cotyledon parenchyma identity. This resource provides a valuable foundation for analyzing gene regulatory programs in soybean cell types across tissues and life stages.

## Introduction

Plants are composed of cells from various tissues and cell types, each containing the same genome, but exhibiting highly divergent gene expression that enables specialized functions. One key driver of transcriptional variation is *cis*-regulatory elements (CREs), non-coding loci in the genome that regulate gene expression in a spatiotemporal manner.^1^ Spatiotemporal gene expression is controlled by interactions between specific binding motif sequences and cognate transcription factors (TFs), along with cofactors assembled at CREs.^2^ Most TFs bind to CREs in nucleosome-depleted accessible chromatin regions (ACRs).^3^ Consequently, distinct TF expression and chromatin accessibility patterns establish the gene expression programs of specific cell types. Thus, detailed maps of CRE accessibility and gene expression in diverse cell types are essential for understanding how different cells use the genome, facilitates our functional understanding of the genome, and enables the exploration of gene regulatory networks.

Advancements in single-cell genomics, such as snRNA-seq (single-nucleus RNA sequencing) and scATAC-seq (single-cell sequencing of assay for transposase accessible chromatin), enable the profiling of transcriptomes and chromatin accessibility from complex tissues at single-cell resolution.^4–6^ Extensive single-cell genomic datasets have been generated by large projects in mammals, such as the Human Cell Atlas and the Mouse Cell Atlas.^7–10^ In plants, single-cell research has mostly been focused on transcriptomes, often limited to selected organs, tissues, and cell types.^11–17^ To date, only three atlas-scale single-cell transcriptomes or chromatin accessibility maps have been reported in *Arabidopsis thaliana*, *Oryza sativa (*rice) and *Zea mays* (maize), each limited to a single modality.^18–20^ However, while extremely valuable, these resources are limited by challenges inherent in single-cell genomic technologies, where the cell types are extracted from their origin in a complex tissue, potentially losing critical biological information, and increasing the difficulty of proper cell-type annotation.^21^

Cell-type annotation is fundamental for elucidating cell population heterogeneity and is typically determined through cell-type markers specifically expressed in one or a few cell types.^12,21^ For many non-model species, there are usually insufficient validated marker genes, and cell-type annotation often relies on the expression patterns of orthologs in model plants, mostly *Arabidopsis*.^14,19^ However, annotation based on ortholog gene expression can be problematic due to gene loss, gene duplication or gene functional diversification following whole genome duplications. Recently, spatial transcriptomics has provided the opportunity to investigate gene expression profiles within the spatial context of cells, successfully assisting cell-type annotations in animals and plants without needing *a priori* cell-type markers.^22–24^ To date, no comprehensive cell-type level atlas has been completed for any plants, which spans gene expression, accessible chromatin regions, and spatially resolved cell-type annotations.

Here, we describe a spatially resolved, multimodal single-cell atlas for the crop species *Glycine max* (soybean), which experienced genome duplications approximately 59 and 13 million years ago, resulting in a highly duplicated genome with nearly 75% of its genes present in multiple copies^25^. We measured chromatin accessibility and gene expression in 316,358 nuclei across ten soybean tissues, which identified and characterized 303,199 ACRs in 103 distinct cell types. We found that nearly 40% of ACRs showed cell-type-specific patterns and were enriched for TF binding motifs controlling cell-type specification and maintenance. Focusing on a unique feature of soybean biology, the infected cells which make up the developing nodules, we identified the non-cell autonomous activity of NLP7 and the conservation of a *NIN* gene regulatory network for legume symbiotic nitrogen fixation. Three sub-cell types of endosperm were detailed characterized and we found that a group of 13 sucrose transporters, including two SWEETs (SUGARS WILL EVENTUALLY BE EXPORTED TRANSPORTERs): *GmSWEET15a* and *GmSWEET10a*, were co-up-regulated in late peripheral endosperm, both sharing the DOF11 binding motif. We also constructed comprehensive developmental trajectories across embryogenesis and early maturation and identified key embryo cell type specification regulators during embryogenesis. Finally, we created an interactive web atlas to disseminate these resources, which we named the soybean multi-omic atlas (https://soybean-atlas.com/).

## Results

### Assembly of a single-cell accessible chromatin and expression atlas in soybean

To generate a comprehensive accessible chromatin and transcriptome atlas across soybean cell types, we collected samples from ten tissues at different stages of the soybean life cycle. These tissues included leaf, hypocotyl, root, nodule, young pod, and five stages of developing seeds: globular stage (GS), heart stage (HS), cotyledon stage (CS), early maturation stage (EMS), and middle maturation stage (MMS). For each tissue, we conducted scATAC-seq and snRNA-seq with at least two replicates, using optimized soybean nuclei isolation methods (Figure 1A, Methods). After filtering out low-quality nuclei and doublets, we obtained high-quality accessible chromatin profiles for ten tissues, totaling 200,732 nuclei with a median of 17,755 unique Tn5 transposase (Tn5) integrations per nucleus, and transcriptome profiles for seven tissues, totaling 115,626 nuclei with a median of 2,474 Unique Molecular Identifiers (UMIs) and 1,986 genes detected per nucleus (Figure S1; Tables S1,2). Initial clustering of 2,000 random nuclei from all tissues revealed similar cluster structures in both scATAC-seq and scRNA-seq, with seed tissue nuclei clearly separated from non-seed tissues (Figure 1B-C). To further explore cell type heterogeneity in soybean tissues, we used the *Seurat*^26^ and *Socrates*^19^ workflows for separate analysis of each tissue. We identified 147 and 97 scATAC-seq and snRNA-seq cell clusters respectively, revealing the diverse cell types or states in soybean (Table S3, 4).

**Figure 1.**
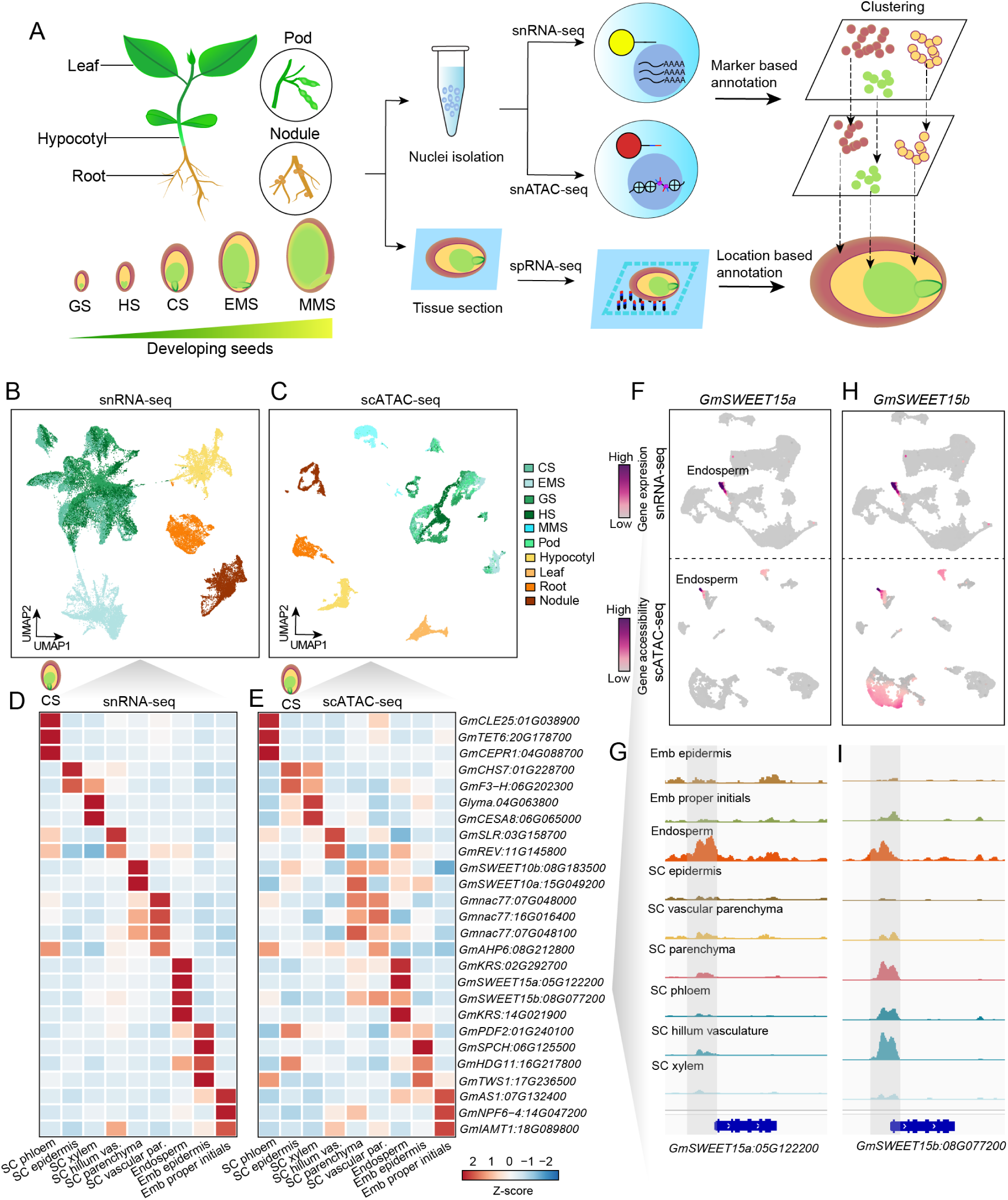
Profiling single-nuclei transcriptomes and chromatin accessibility in soybean. **(A)** Overview of tissue types and experimental design. Seed stages include GS (globular stage), HS (heart stage), CS (cotyledon stage), EMS (early maturation stage), and MMS (middle maturation stage). **(B-C)** Two-dimensional embeddings using Uniform Manifold Approximation and Projection (UMAP) depicting similarity among nuclei based on gene expression (B) and gene chromatin accessibility (C). 2,000 nuclei were randomly selected from each tissue and colored by tissue type. **(D-E)** Z-score heatmap of gene expression (D) and gene chromatin accessibility (E) for representative marker genes across shared cell types in soybean cotyledon stage seeds. SC, seed coat; Emb, embryo. **(F-G)** UMAP embeddings overlaid with gene expression (**top**) or gene accessibility (**bottom**) (F) and pseudobulk cell type Tn5 integration site coverage (G) around the endoderm marker gene *GmSWEET15a*. **(H-I)** Similar to panels F-G, but for the paralog gene *GmSWEET15b*.

To annotate these cell clusters, we collected a set of marker genes from the literature spanning multiple species, including soybean, *Arabidopsis*, and maize, and matched them to expected soybean cell types. Cell types were assigned based on a manual review of marker gene performance and evaluation of enriched biological processes (Methods, Table S5). For example, in cotyledon stage seeds, we identified 17 clusters in scATAC-seq and 18 clusters in snRNA-seq, with high concordance between the two replicates (Figure S2A-D). By comparing the single-cell data with previously published laser capture microdissection RNA-seq datasets^27,28^, we identified the three main regions of soybean seeds: seed coat, endosperm, and embryo, as well as specific cell types, such as the seed coat endothelium and seed coat inner integument (Figure S2E, F). Additional cell types were annotated based on representative marker genes. For instance, the plasma membrane sugar transporter GmSWEET15, which mediates sucrose export from the endosperm to the embryo.^29^ As expected, the paralogs *GmSWEET15a* and *GmSWEET15b* showed both expression and chromatin accessibility enriched in the endosperm, with neighboring ACRs reflecting the potential *cis*-regulatory elements driving its endosperm specific gene expression (Figure 1F-I). After comprehensive annotation and subsequent analysis, we identified a total of 103 and 79 cell types in the scATAC-seq and snRNA-seq data, respectively, with a high correlation between gene accessibility from scATAC-seq and gene expression from snRNA-seq for the same cell types (Figure S3-6, Table S3-5).

### Validation of cell-type identity with spatial transcriptomics

The limited availability of experimentally validated marker genes for cell-type annotation in scATAC-seq and scRNA-seq datasets is a common challenge, particularly in non-model species. Homology-based marker identification is problematic due to gene loss, duplication, or neofunctionalization. To validate the cell-type annotations for the single-cell datasets, we conducted spatial RNA-seq (spRNA-seq) for five tissue types matching the single-cell datasets (root, hypocotyl, seed at heart stage, cotyledon stage, and early maturation stage). Multiple serial tissue sections were placed on a 10X Genomics Visium spatial slide. In total, we profiled 12,490 high-quality spatial spots across these tissues (Table S6). The median gene number per spot ranged from 453 to 6,262 across all tissue types.

The unsupervised clustering of the expression profiles revealed that spatial spot clusters showed cell-type specific spatial localization (Figure 2B and Figure S7B). For example, we identified 13 unique clusters in the cotyledon stage seed dataset (Figure 2B). Four of these clusters are localized in the embryo region, three in the endosperm region, and six within the seed coat region (Figure 2B). This indicates high-quality spatial transcriptome data and enables us to accurately annotate cell types based on tissue histology. The Visium spatial slides are designed with 55-um resolution spots, which capture gene expression profiles from multiple cells. To study the spatial expression profile at single-cell resolution and validate the snRNA-seq cell-type annotation, we performed the deconvolution analysis using spRNA-seq and snRNA-seq datasets of the same tissue types. The prediction score of each snRNA-seq cell was calculated to quantify the certainty of the association between snRNA-seq cells and their predicted spatial spots. We observed high prediction scores between similar cell types that were independently annotated in the two datasets (Figure 2C, Figure S7C), supporting a robust annotation.

**Figure 2.**
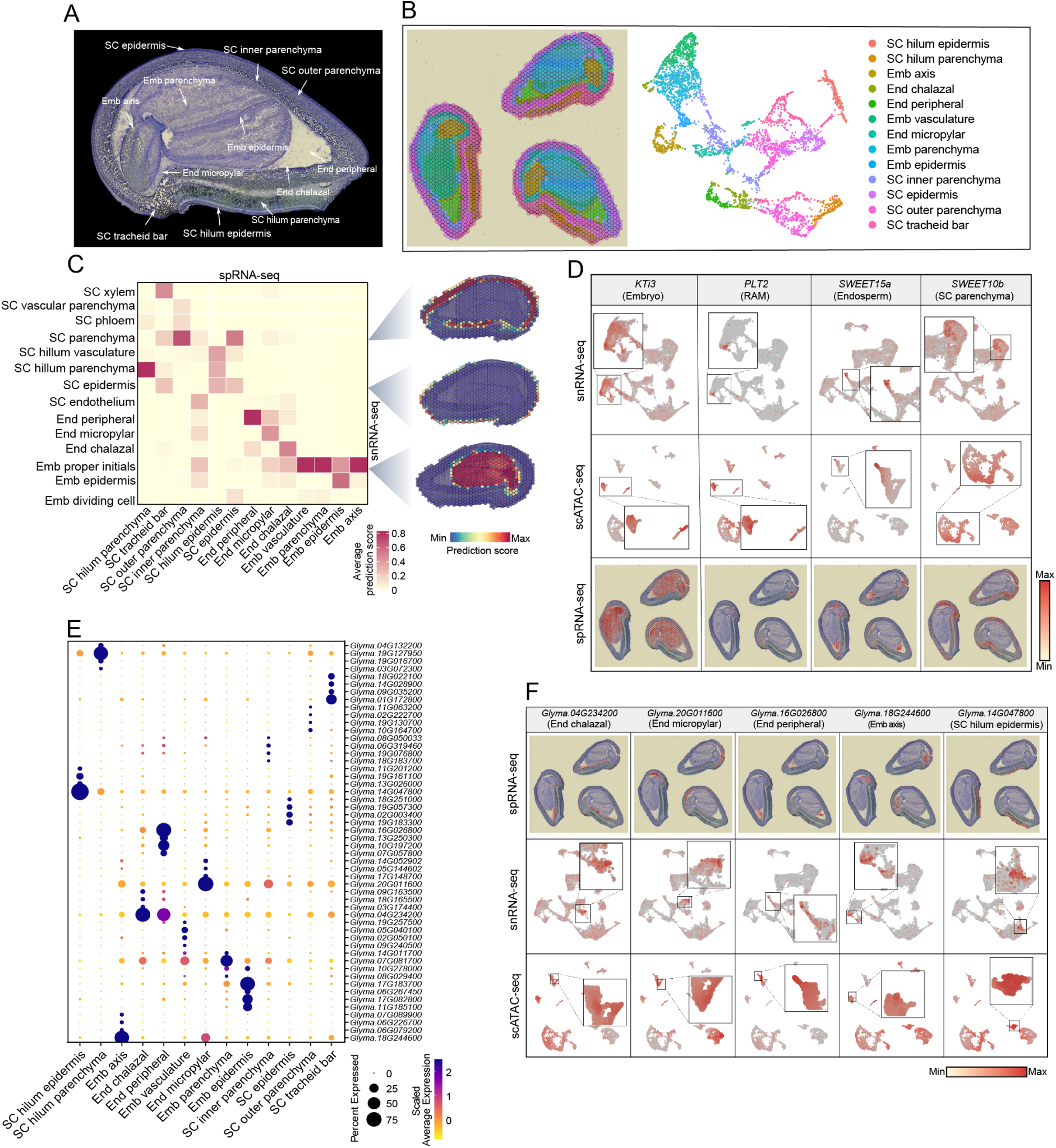
A spatially resolved transcriptome facilitates cell-type annotation for soybean seeds. **(A)** The histological structure of soybean seeds at the cotyledon stage. **(B)** The visualization of spatial spot clusters on the tissue section (left) and on the UMAP plot (right). **(C)** Heatmap of the snRNA-seq cell-type prediction scores on the spRNA-seq cell types (left) and the spatial distribution of predicted snRNA-seq cell types on the tissue section (right). **(D)** The validation of known marker genes used in the scRNA-seq data. The gene expression of selected markers was plotted on the UMAP of snRNA-seq data (top), scATAC-seq data (middle), and on the spatial plot of the tissue section (bottom). **(E)** Dotplot of the top *de novo* marker genes identified for each cell type in the spRNA-seq data. **(F)** The validation of spatial *de novo* marker genes in the single-cell data. The gene expression of selected markers was plotted on the spatial plot of the tissue section (top), the UMAP of snRNA-seq data (middle), and the scATAC-seq data (bottom).

Leveraging the spatial transcriptome data, we corroborated the known marker genes selected for the snRNA-seq cell-type annotation (Figure 2D and Table S5). For example, *GmKTi3* (*Glyma.08G341500*) mRNA is known to be exclusive to the soybean embryo,^30^ and we confirmed *GmKTi3* embryo specificity with the spRNA-seq data. Likewise, *PLETHORA2* (*PLT2*) is expressed in the *Arabidopsis* root apical meristem (RAM)^31^, which was validated by the spatial transcriptomic data. Finally, *GmSWEET15a* is mainly expressed in the cotyledon stage endosperm, which is also consistent with our spRNA-seq data; the seed coat parenchyma marker *GmSWEET10b* (*Glyma.08G183500*)^32^ showed a highly specific expression in the seed coat. Collectively, these data support that the spRNA-seq results accurately reflect mRNA localization and provide a valuable tool for marker *in situ* validation.

To identify more soybean cell-type-specific markers, we performed *de novo* marker identification using the spRNA-seq and snRNA-seq datasets (Figure 2E and Figure 2F, Table S7, S8). With the *de novo* markers from spRNA-seq, we distinguished similar cell types that are spatially differentiated. For example, we identified three subclusters of endosperm cells, and annotated them as micropylar, peripheral, and chalazal endosperm based on their localization in the seed (Figure 2F). The spatial *de novo* markers from these cell types showed distinct expression patterns in the corresponding snRNA-seq and scATAC-seq subclusters. Taken together, by integrating the spRNA-seq, we not only validated the cell-type annotation for snRNA-seq and scATAC-seq, but also identified spatially differentiated sub-cell types of endosperm.

### Identification and characterization of ACRs across cell types

To identify ACRs in the 103 cell types, we aggregated chromatin accessibility profiles from all nuclei within each cell cluster and applied a peak calling procedure optimized for single-cell data (Methods). This uncovered 303,199 non-overlapping ACRs, ranging from 137,046 to 193,792 per tissue (Figure 3A). Compared to bulk ATAC-seq from leaf at the same stage (Methods), scATAC-seq identified almost twice as many ACRs despite having fewer total reads, as scATAC-seq identified cell-type-specific ACRs (Figure 3B, C). Next, we categorized the ACRs based on their proximity to annotated genes: 128,916 (45.52%) overlapped genes (genic ACRs), 74,655 (24.62%) were within 2 kilobases (kb) of genes (proximal ACRs), and 99,628 (32.86%) were more than 2 kb away from genes (distal ACRs). Distal ACRs had significantly higher cell-type specificity scores than genic ACRs and proximal ACRs, suggesting their important role in establishing cell-type-specific gene expression patterns (t.test, p-value < 2.2e^-16^, Figure S8A). Genetic diversity from the soybean haplotype map (GmHapMap)^33^ was remarkably reduced, and TF motifs were enriched at the summit of all three groups of ACRs, supporting the functionality of the identified ACRs (Figure 3E, Figure S8B).

**Figure 3.**
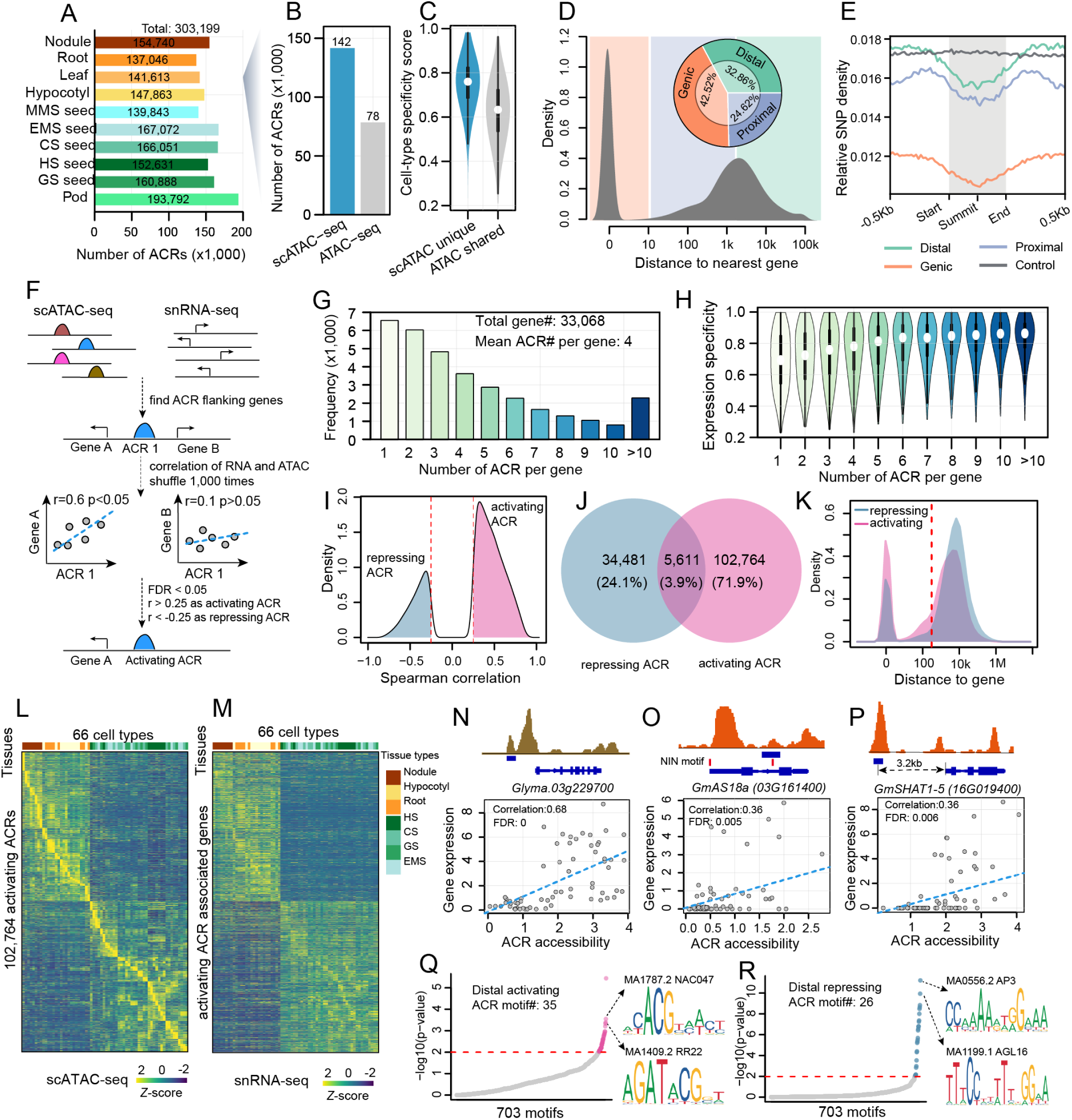
Characterization of ACRs across cell types. **(A)** Number of ACRs identified in each tissue. **(B)** Comparison of the number of ACRs identified using scATAC-seq versus bulk ATAC-seq in leaf tissues. **(C)** Distribution of cell-type specificity score for ACRs shared between bulk ATAC and scATAC, and those unique to scATAC-seq. **(D)** Bimodal distribution of ACR distances to the nearest gene. ACRs are categorized into three groups based on the distance from the summit to the nearest gene: genic ACRs (overlapping or within 10 bp of genes), proximal ACRs (within 2 kb of genes), and distal ACRs (more than 2 kb away from genes). **(E)** Relative SNP density within 500-bp flanking regions of different classes of ACRs and control regions. **(F)** Schematic overview of the computational strategy used to predict the activity function of ACRs. **(G)** Distribution of genes associated with different numbers of ACRs. **(H)** Distribution of expression specificity for genes associated with different numbers of ACRs. **(I)** Density distribution of the overall Spearman correlation coefficient between ACRs and flanking genes. **(J)** Venn diagram analysis of activating and repressing ACRs. **(K)** Density distribution of the distance between the pair of ACRs and genes for the activating and repressing ACRs. **(L-M)** Heatmap showing chromatin accessibility of activating ACR (L) and the expression of associated genes (M). **(N-P)** Pseudobulk cell type Tn5 integration site coverage patterns around gene bodies (top) and scatter plots of ACR accessibility and gene expression across 66 cell types (bottom) for *Glyma.03g229700*, *GmAS18a (03G161400)*, and *GmSHAT1-5 (16G019400)*, respectively. (Q-R) TF motif enrichment of distal activating ACRs (Q) and distal repressing ACRs (R).

ACRs can be classified as activating ACRs, which positively regulate gene expression, and repressing ACRs, which reduce gene expression.^34^ To predict ACR function, we associated ACRs with putative target genes based on the correlation between ACR accessibility and nearby gene expression across all cell types in the scATAC-seq and snRNA-seq datasets (Figure 3F, Methods). This process identified 145,638 ACR-gene associations for 137,245 ACRs and 33,068 genes, with an average of four ACRs per gene (Figure 3G, Table S9). We found that gene expression cell-type specificity is positively correlated with the number of associated ACRs, suggesting that the number of ACRs is associated with restricted gene expression patterns (Figure 3H). Next, we categorized ACRs with positive correlations as activating ACRs and those with negative correlation as repressive ACRs (Figure 3F, I, L, M; Figure S8C). Overall, 71.9% were activating ACRs, 24.1% were repressing ACRs, and 3.9% had ambiguous functions with mixed significant positive and negative correlations with flanking genes (Figure 3J). Activating ACRs were more likely to act proximally compared to repressing ACRs (Figure 3K). Notably, we identified three known activating CREs expressed in different tissues and involved diverse developmental pathways (Figure 3N-P), such as in seed tissues,^35^ *ASYMMETRIC LEAVES2-LIKE 18 (ASL18)*, a known root nodule symbiosis marker,^36^ and a pod shattering-resistance related gene^37^.

To identify motifs that could act as distal activators or repressors, we conducted a TF motif enrichment analysis on the distal activating and repressing ACRs. We found 35 motifs enriched in distal activating ACRs, and six of the top ten motifs had known transcriptional activator activity, such as NAC DOMAIN CONTAINING PROTEIN 47 (NAC047)^38^ and RESPONSE REGULATOR 22 (RR22)^39^ (Figure 3Q, Table S10). Additionally, 26 motifs were enriched in distal repressing ACRs, primarily Type II MADS-box factors like APETALA3 (AP3)^40^ and AGAMOUS-LIKE 16 (AGL16)^41^, known transcriptional repressors involved in floral organ specification (Figure 3R, Table S10). Type II classic MADS-box genes are key developmental regulators in angiosperms and are well-studied due to their role in floral organ specification.^42^ We observed distinct MADS gene expression patterns in seed versus non-seed tissues, consistent with MADS-box genes regulating reproductive growth by transcriptionally repressing distal genes. In summary, we constructed a comprehensive atlas of *cis*-regulatory activity across 103 soybean cell types, predicted their target genes and regulatory functions by integrating snRNA-seq data. These results provide a foundation for dissecting gene regulatory programs at cell-type resolution.

### Identification and characterization of cell-type-specific ACRs (ctACRs)

This single-cell atlas provides an excellent opportunity to characterize the heterogeneous regulatory programs underlying specialized cell-type functions. First, we identified ctACRs that were significantly more accessible in one or two cell types within each tissue (Methods). Approximately 40.23% of the ACRs (122,558 ACRs) were identified as ctACRs across ten tissues, ranging from 12,711 in root to 37,897 in young pod (Figure 4A, Figure S9A, Table S11). We observed a higher number of ctACRs in seed-related tissues compared to non-seed tissues, with a significantly higher number of endosperm-specific ACRs in young developing seeds compared to the ctACR number in other cell types (Figure S9A). The proportion of ACRs located in proximal regions was similar across ctACRs and non-ctACR, but there was a higher proportion of distal ACRs among ctACRs (Figure S9C). This suggests the importance of distal ACRs in contributing to cell-type-specific chromatin accessibility patterns. Comparing polymorphism density across distal specificity groups, we found that ctACRs were highly conserved, suggesting positive selection of ctACRs in soybean breeding (Figure S9D).

**Figure 4.**
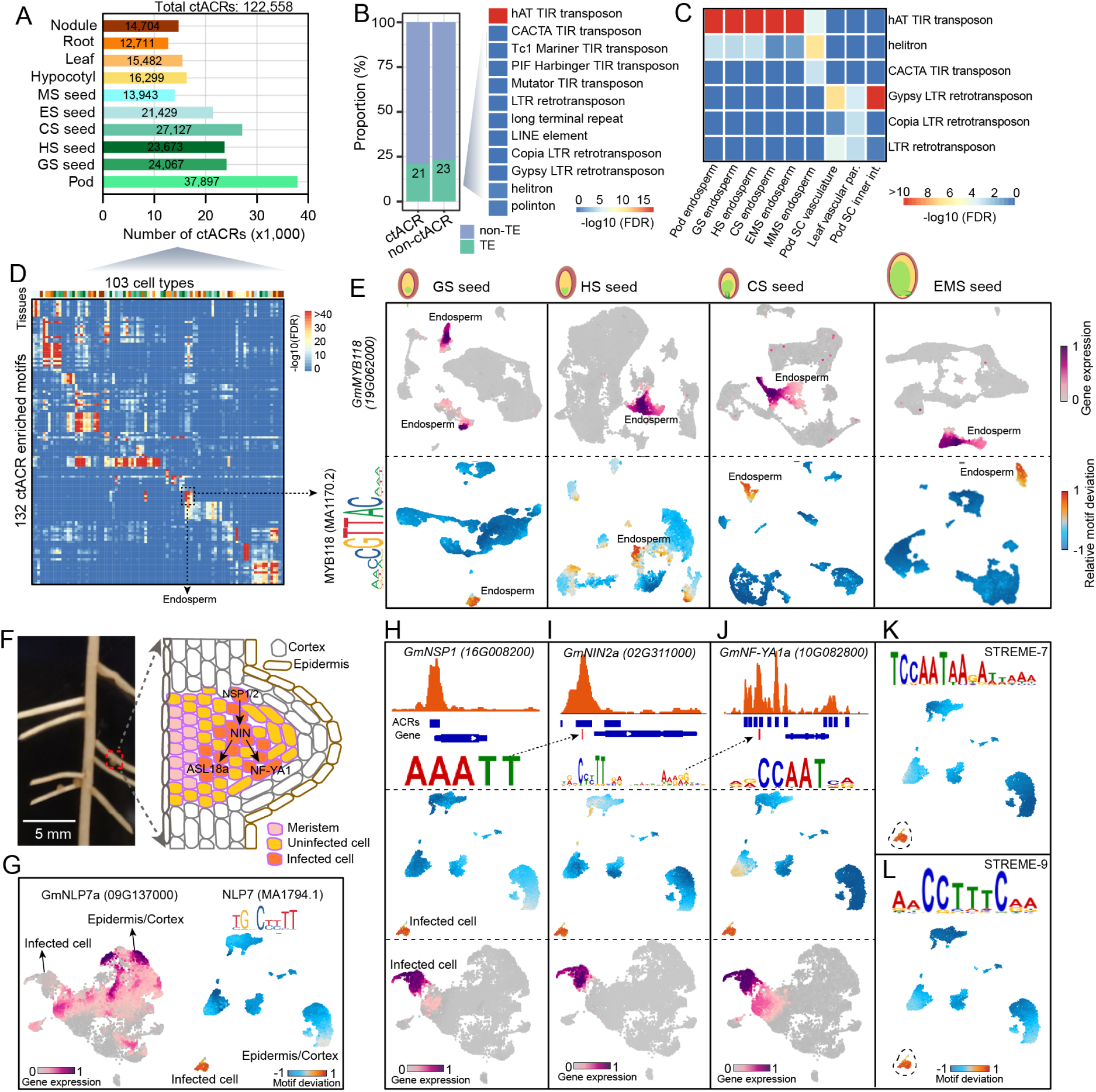
Characterization of cell-type-specific ACRs, motif and TFs. **(A)** Number of ctACRs identified in each tissue. **(B)** Proportion of ACRs that overlap with TEs and TE enrichment in all ctACRs. **(C)** TE enrichment in ctACRs for each cell type. **(D)** Heatmap of TF motif enrichment across 103 cell types. **(E)** UMAP embeddings overlaid with gene expression of *GmMYB118* (top row) or TF motif deviation score of the MYB118 binding motif (bottom row) across four developmental stages of seeds. **(F)** Image of a root with nodules (left) and an illustration of major cell types and the gene regulatory pathway in infected cells of developing nodules. **(G)** UMAP embeddings overlaid with gene expression of *GmNLP7a* and TF motif deviation score of NLP7 in nodule tissue. **(H-J)** Pseudobulk cell type Tn5 integration site coverage pattern around gene body (top), UMAP embedding overlaid motif deviation score (middle) and gene expression (bottom) for *GmNSP1*(H), *GmNIN2a* (I) and *GmNF-YA1a* (J). **(K-L)** UMAP embedding overlaid TF motif deviation score for *de novo* motifs of STREME-7 and STREME-9

Transposable elements (TEs) contribute to cell-type-specific CREs in both mammals and plants.^19,43,44^ For example, enhancer cell-type-specific CREs are often found within long terminal repeat retrotransposons (LTRs) in maize.^19^ In soybean, a similar proportion of ctACRs and non-ctACRs overlapped with TEs. TE enrichment analysis indicated significant enrichment of hAT TIR transposons in ctACRs (Fisher’s exact test, FDR < 10e^-16^), representing a distinct TE family enrichment as compared to maize. To investigate the role of TEs and their relationship to cell-type-specific CREs, we conducted an enrichment analysis comparing ctACRs-overlapping TEs with non-ctACRs-overlapping TEs for each cell type. We found significant TE enrichment in nine cell-type states (Fisher’s exact test, FDR < 0.01). Notably, hAT TIR transposons were significantly enriched in endosperm-specific ACRs across all seed development stages (FDR < 10e^-4^, Figure 4C), highlighting a unique relationship between a specific TE family and cell-type critical for agriculture.

### Identification of key TF regulators that define distinct cell identities

Identifying which TFs are involved in generating and maintaining a diversity of cell types from an invariant genome is a central question in developmental biology. We leveraged these data to systematically assess which TF motifs are enriched in ctACRs across tissues, thus identifying key regulatory networks potentially critical in cell fate specification.

Initially, for each cell type, we determined (Fisher’s Exact test) which TF motifs are overrepresented in ctACRs compared to non-ctACRs. By analyzing each tissue independently, we identified the most highly enriched TF motifs and TFs from the JASPAR database^45^ for 103 cell types across all tissues, revealing both known and novel potential regulators (Figure 4D, Figure S9E, Table S12). For example, the HDG11 (MA0990.2) motif, an established regulator of epidermal cells^46^, is highly accessible in epidermal cells of hypocotyl, root, leaf, and cotyledon stage seeds. It is likely that HDG11 and its family members are critical drivers of epidermal cell fate. Similarly, the DOF1.6 (MA1275.1) motif is enriched in procambium-related cells across all tissues (Figure S9F, Table S12). Additionally, the MYB118 motif, a known endosperm-specific transcriptional activator^47^, is enriched for cell-type-specific chromatin accessibility in endosperm and is specifically expressed in soybean endosperm cells across four developmental stages (Figure 4E, Figure S9F). These results show that specific TF motifs and their associated networks are used in a tissue-specific and cell-type-specific manner.

Adapting these analyses, we were further interested in developing nodules, where a symbiosis between legumes and soil bacteria fix nitrogen for both the plant and the natural or agricultural ecosystem.^48^ Nitrogen fixation occurs in infected cells, a unique cell type that encapsulates the bacteria (Figure 4F). However, how these cells are altered in terms of their CRE usage after infection remains underexplored. We found a series of symbiotic nitrogen fixation genes that were specifically expressed and accessible in these infected cells in both snRNA-seq and scATAC-seq datasets (Figure S3C, D). 73 TF motifs were enriched in infected cells, including the binding motif of NIN-LIKE PROTEIN 7 (NLP7), a known regulator of root nodule symbiosis^49,50^ (Figure 4G, Table S12). Notably, there was a spatial separation between *NLP7’s* expression in epidermis or cortex and its binding site accessibility in infected cells, suggesting non-cell autonomous activity, following a previously published method for identifying non-cell autonomous TFs^19^ (Figure 4G). The top two most enriched motifs in infected-cell-specific ACRs were AHL13 (MA2374.1), which regulates jasmonic acid biosynthesis and signaling^51^ and ANTHOCYANINLESS 2 (MA1375.2) which regulated anthocyanin accumulation and primary root organization^52^ (Figure S9G, H).

Only seven of the motifs in the JASPAR database^45^ are from soybean, with most being from *Arabidopsis* (580) or other species (218), potentially limiting the study key soybean TF motifs, as they are unknown. For example, key regulator genes essential for initiating cortical cell divisions and microbial infection during nodulation, such as NODULATION SIGNALING PATHWAY 1 (NSP1)^36^, NODULE INCEPTION (NIN)^36^, ASYMMETRIC LEAVES 2-LIKE 18 (ASL18)^36^, Nuclear Factor-YA1 (NF-YA1)^48^, were highly expressed in infected cells (Figure 4H-J). Their TF binding motifs, characterized in *Medicago truncatula* and *Lotus japonicus*, were expected to be enriched in infected-cells-specific ACRs, but they were absent in the JASPAR database. Using the same analysis, we found those TF motifs were enriched and showed specific chromatin accessibility in infected cells, suggesting their conservation in soybean (Figure 4H-J, Table S12).

To comprehensively identify potential TF binding motifs in infected cells, we performed *de novo* motif enrichment in infected-cell-specific ACRs, identifying 10 enriched motif clusters (Table S13). Interestingly, all four binding motifs of known key regulators (NLP7, NIN, NSP1, NF-YA1) matched the *de novo* motifs (Figure S9I). Additional TF motifs matched known motifs in the JASPAR database, including binding sites for AP2/ERFs, B3 domain-containing TFs RAV2, Basic leucine zipper (bZIP) TFs, Ethylene-responsive (ERF) TFs, and Protein BASIC PENTACYSTEINE1 (BPC1) TFs. Notably, among these motifs, the GCC-box motif is a known pathogenesis-related promoter element that recruits ERF TFs, including the Ethylene Response Factor Required for Nodulation1 (ERN1), which is essential for infection-thread formation and nodule organogenesis in *Medicago*.^53^ We also identified two novel motifs, which are specifically accessible in the infected cell, including the AACCTTTCAA motif (STREME-7) and the TCCAATAAGATTAAA motif (STREME-9) (Figure 4K, L), which suggests their importance for nodule development in soybean and provides clues into uncharacterized nodulation transcriptional regulatory circuits. In summary, integrating TF motif enrichment in ctACRs with scRNA-seq allows us to profile known TF binding motifs of key regulators and *de novo* uncover novel TF motifs essential for cell-type specification.

### Characterizing three sub-cell types of endosperm across seed development

The endosperm plays a crucial role in supporting embryo growth by supplying nutrients and other factors during seed development.^54–56^ Soybean endosperm is a membrane-like, semi-transparent tissue between embryo and seed coat. Primary endosperm can be divided into three sub-cell types: micropylar, nearest to the young embryo; peripheral, in the center of the endosperm region; and chalazal, at the opposite end of the embryonic axis, towards the seed coat attachment point (Figure 5A).^57^ Although the development of these subregions has been well-characterized morphologically, little is known about the molecular processes occurring in these subregions or how their development is coordinated within the context of seed maturation.

**Figure 5.**
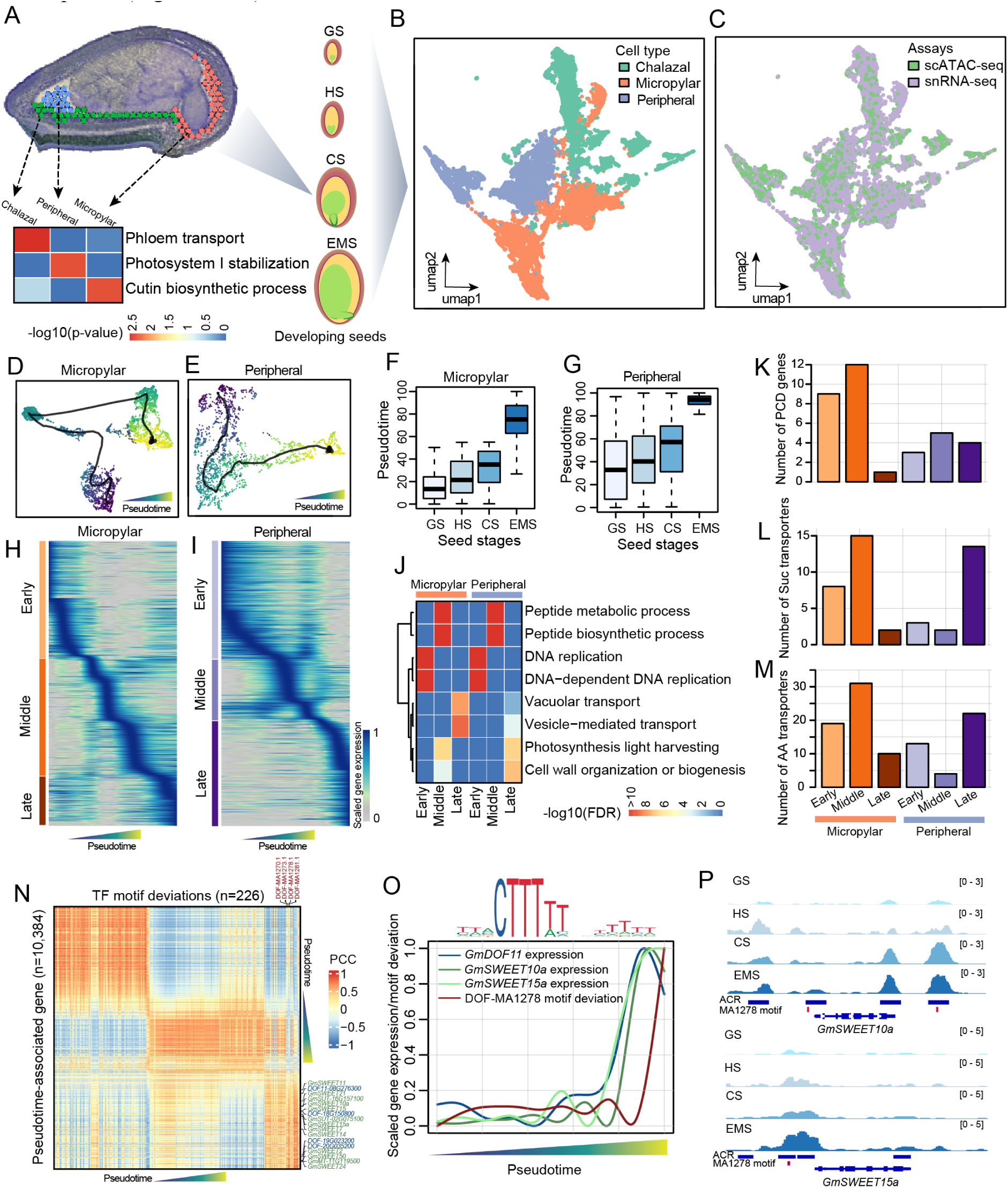
Characterizing three endosperm sub-cell types across seed development. **(A)** Spatial tissue section showing the three sub-cell types (chalazal, peripheral, micropylar endosperm) (top) and a heatmap of their representative enriched biological processes (bottom). **(B-C)** UMAP embeddings overlaid with cell type (B) or assays (C) **(D-E)** UMAP embeddings depicting pseudotime trajectories for micropylar endosperm (D) and peripheral endosperm (E). **(F-G)** Comparison of pseudotime and categorical seed stages for micropylar endosperm (D) and peripheral endosperm (E). **(H-I)** Heatmap of pseudotime-associated genes (FDR < 0.05) for micropylar endosperm (H) and peripheral endosperm (I). **(J)** Heatmap of representative enriched biological processes across pseudotime-inferred stages and cell types. **(K-M)** Number of programmed cell death genes (K), sucrose transporters (L), and amino acids transporters (M) across pseudotime-inferred stages and cell types. **(N)** Correlation heatmap between TF motif deviation scores and pseudotime-associated genes aligned by pseudotime for peripheral endosperm. PCC, Pearson correlation coefficient **(O)** Expression of *GmDOF11 (08G276300)*, DOF-MA1278 motif deviation, and expression of its putative target genes *GmSWEET10a* and *GmSWEET15a*. The DOF-MA1278 motif is shown above. **(P)** Pseudobulk cell type Tn5 integration site coverage around *GmSWEET10a* and *GmSWEET15a* across the four seed stages.

By integrating snRNA-seq and spatial RNA-seq, we separated the three sub-cell types of endosperm (Figure 2B) and gained insights into the cellular processes within each sub-cell type by identifying significantly overrepresented Gene Ontology (GO) terms (P < 0.01, Figure 5A, Table S14). Some of overrepresented GO terms were consistent with the known roles of these endosperm sub-cell types in seed development. For example, the peripheral endosperm is enriched in photosynthesis-related pathways, consistent with the presence of chloroplasts^57,58^, the chalazal endosperm is enriched in vascular transport pathways, aligning with its role in loading maternal resources into developing seeds^56,59^, and the micropylar endosperm is enriched in cutin biosynthetic process pathways, suggesting involvement in cuticle synthesis in the nearby embryo epidermis^54,55,60^. These results support the reliability of the annotation of the three sub-cell types of endosperm cells.

To overview endosperm development, we analyzed all endosperm nuclei across four stages (globular, heart, cotyledon, and early maturation) of seed development, integrating scATAC-seq and snRNA-seq modalities (Figure 5B-C, Figure S10A, Methods). Using *de novo* markers from spRNA-seq, we clearly separated and annotated the three sub-cell types (Figure S10B, Table S15). Comparing the proportion of nuclei in each stage across clusters revealed a developmental change in cell number for peripheral and micropylar endosperm, but not for chalazal endosperm (Figure S10C-H). This observation can be explained by the cellularization of peripheral and micropylar endosperm following nuclei proliferation, while the chalazal endosperm undergoes degradation without a clear cellularization process.^57,59^

To determine regulatory and gene expression dynamics during endosperm development, we performed pseudotime analysis for micropylar and peripheral endosperm using snRNA-seq nuclei as a reference (Figure 5D, E). Pseudotime was highly correlated with the progressive development (Figure 5F, G). We classified genes based on expression patterns across pseudotime into three stages (early, middle, late) for micropylar and peripheral endosperm (Figure 5H, I, Table S16,17). GO enrichment analysis reflected the processes of nuclei proliferation in the early stage and further cellularization and function specification in later stages (Figure 5J, Table S18). These results suggested we constructed a comprehensive developmental trajectory for micropylar and peripheral endosperm, allowing high resolution exploration of the gene regulatory network along the endosperm development.

During soybean seed development, endosperm cells undergo programmed cell death (PCD) and transfers nutrients to support rapid embryo growth and expansion.^56,61,62^ The molecular regulation of endosperm PCD, and which nutrient transporters are involved, remains poorly understood. By examining expression patterns of PCD-related genes^63^ and sucrose or amino acid transporter genes^64^ in developmental trajectories, we found more PCD-related and nutrient transporter genes expressed in early and middle stages of micropylar endosperm than the late stage (Figure 5K-M, Table S16,17). The micropylar endosperm, being closest to the embryo, undergoes PCD and serves as an important nutrient source during early seed development.^61^ More nutrient transporter genes were expressed in the peripheral endosperm in the late stage, suggesting its role in transferring maternal nutrients in later embryo development.

Sucrose is the major photosynthetic product transported into seeds^65^ and sugar transporters essential for embryo development have been identified and characterized in different plants.^66^ We identified a cluster of 13 sugar transporters highly upregulated in the late stage of peripheral endosperm, including *GmSWEET10a* and *GmSWEET15a*, known to control soybean seed size and oil content^29,32^. As these sugar transporters share similar expression patterns along development, we hypothesize they might share similar TF motif sequences and chromatin accessibility patterns and be regulated by TFs with colocalized expression patterns. To predict shared upstream regulators controlling the 13 sucrose transporters, we scanned all TF motifs in their proximal and genic ACRs, and found five motifs from three TF superfamilies shared by all ACRs (Figure S10I-K): DOF (DNA binding with one finger) family, Homeodomain-leucine zipper (HD-Zip) TFs, and C2H2 zinc-finger TFs, including INDETERMINATE DOMAIN (IDD) TFs. We imputed TF motif deviations from scATAC-seq onto snRNA-seq nuclei, identifying 226 TF motifs following the trajectory pattern, with only two DOF motifs highly correlated with the 13 sugar transporter genes (Figure 5N, Figure S10I-K). We identified four DOF genes highly expressed in the late stage of peripheral endosperm, including *GmDOF11a* (*Glyma.08G276300*), whose paralog *GmDOF11b* (*Glyma.13G329000*) controls soybean seed size and oil content.^67^ Specifically, *GmSWEET10a* and *GmSWEET15a* were highly expressed in the late stage, and their ACRs capturing DOF motif (MA1278), become more accessible throughout seed development (Figure 5O, P).

### Developmental trajectories defining soybean embryogenesis

Many important soybean agronomic traits are established during early seed development. However, the regulatory and gene expression dynamics underlying cellular diversification during embryogenesis and the relationship with agronomically important traits are unresolved. Motivated by this question, we isolated all embryo-related nuclei across four stages (globular, heart, cotyledon, and early maturation) of seed development and performed an integration across scATAC-seq and snRNA-seq modalities (Figure S11A-S11B, Table S19). To improve the resolution of developmental progression, we inferred the precise developmental age of each nucleus using a recently described LASSO regression approach (Figure 6A).^68^ The predicted continuous developmental ages from the full data set (Pearson’s correlation = 0.93) and withheld test nuclei (Pearson’s correlation = 0.96) were highly correlated with the known seed stage (Figure 6B, Figure S11C). We identified 248 genes predictive of developmental age and uncovered the sequential gene expression dynamics associated with overall developmental progression regardless of cell lineage (Figure 6C-6D). These results provide a useful benchmark for anchoring analyses of cellular diversification during embryogenesis.

**Figure 6.**
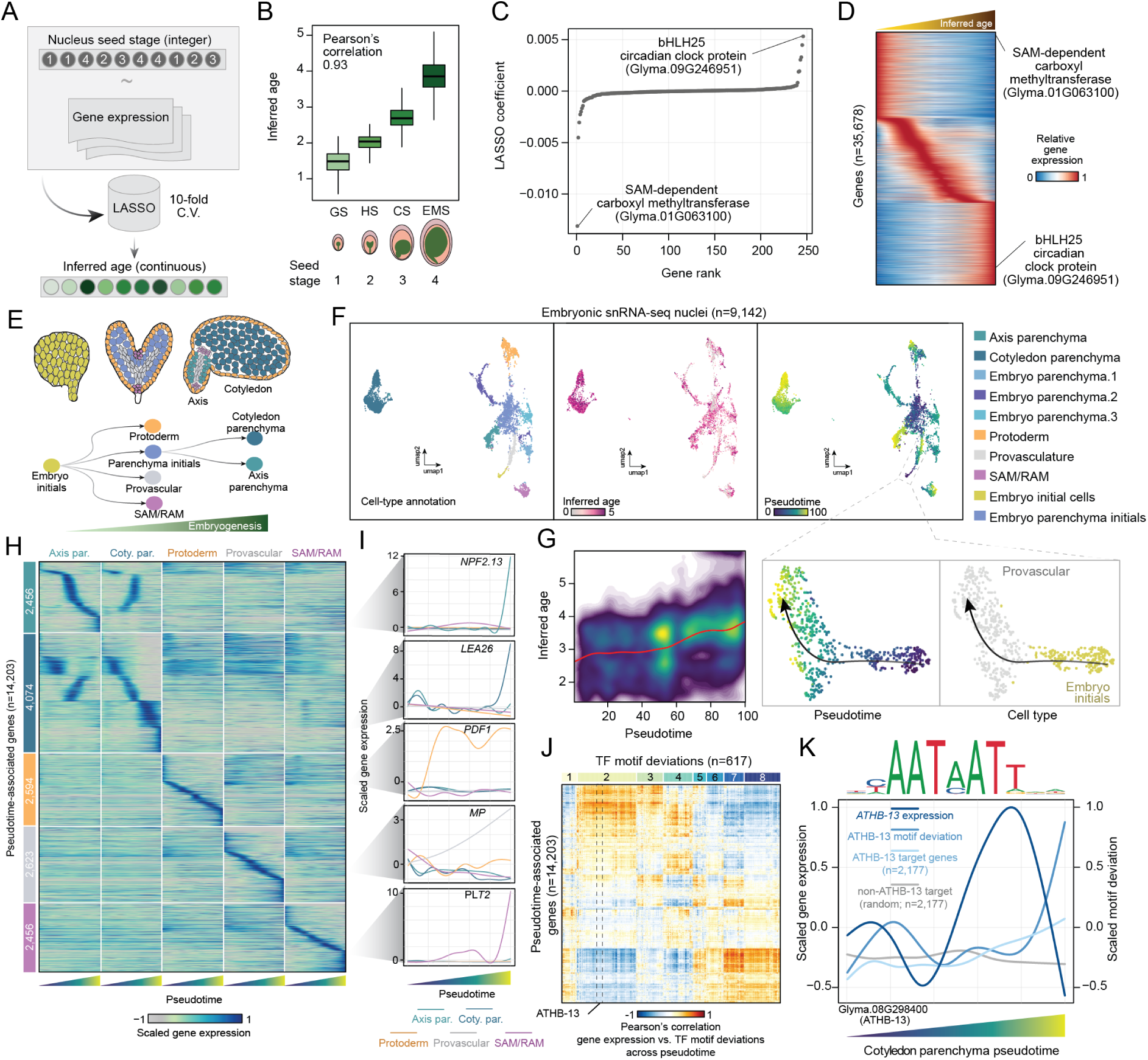
Developmental trajectories defining soybean embryogenesis. **(A)** Illustration of LASSO models to learn continuous representations of nuclei age. **(B)** Comparison of inferred nuclei age and categorical seed stages. **(C)** LASSO coefficient ranks of genes towards inferred nuclei ages. **(D)** Heatmap of relative gene expression levels ordered by nuclei age. **(E)** Schematic of embryogenesis trajectories. **(F)** UMAP scatter plots of cell-type annotation (left), inferred nuclei age (middle), and pseudotime (right) for embryonic snRNA-seq nuclei. **(G)** Comparison of inferred age and pseudotime scores across all embryonic nuclei. **(H)** Heatmap of pseudotime-associated genes (FDR < 0.05) for all five trajectories. **(I)** Exemplary gene expression profiles across pseudotime for five marker genes. **(J)** Correlation heatmap between TF motif deviation scores and pseudotime-associated genes aligned by cotyledon parenchyma pseudotime. **(K)** *ATHB-13* gene expression, ATHB-13 motif deviation, ATHB-13 target gene expression, and control gene expression profiles across cotyledon parenchyma developmental pseudotime. The motif recognized by ATHB-13 is shown above.

Evaluation of cellular diversity across the four seed stages of embryogenesis revealed five distinct developmental branches (Figure 6E). To determine the regulatory and gene expression dynamics that make these lineages unique, we constructed pseudotime trajectories for each individual branch using the snRNA-seq nuclei as a reference. Providing a firm biological foundation, we observed a strong positive trend between pseudotime scores and inferred developmental age (Figure 6F-6G). Interestingly, we found a strong negative correlation between transcriptional complexity and inferred developmental age, a notable feature of differentiation in mammals^68^ that appears to be conserved in plants (Figure S11D). Hypothesizing that cellular diversification would be accompanied by the acquisition of specialized gene expression programs, we identified differentially expressed genes across pseudotime for each individual branch. Indeed, visualization of pseudotime-associated genes revealed unique transcription dynamics for each lineage (Figure 6H). Importantly, we found that several well-known marker genes displayed expected developmental transcription patterns, including *LATE EMBRYOGENESIS ABUNDANT 26* (*LEA26*) in cotyledon parenchyma^69^, *PROTODERMAL FACTOR 1* (*PDF1*) in the protoderm, *MONOPTEROS* (*MP*) in provascular, and *PLETHORA2* (*PLT2*) in shoot/root apical meristem trajectories that collective support robust trajectory ordering (Figure 6I).^70^

Specification of the developing cotyledon parenchyma has been posited as a key developmental stage that determines nutrient composition of mature seeds (Figure 6E).^71^ We hypothesized that detailed interrogation of the regulatory dynamics between cotyledon and axis parenchyma would be informative for understanding the divergence of these tissues during embryogenesis and uncover ideal targets for soybean improvement efforts. To this end, we imputed TF motif deviations from scATAC-seq onto embedded snRNA-seq nuclei (Figure S11E, S6H) and identified TF motif deviations and gene expression patterns that were correlated across pseudotime for the cotyledon parenchyma trajectory (Figure 6J). This analysis revealed eight TF modules associated with largely distinct gene sets, representing putative gene regulatory networks underlying cotyledon parenchyma development. Next, we speculated that temporal gene expression divergence between axis and cotyledon parenchyma could identify genes associated with lineage bifurcation of parenchyma initials. By comparing temporal gene expression between axis and cotyledon parenchyma, using each branch as a reference (Figure S11F), we found similar gene expression patterns between axis and cotyledon parenchyma early in both trajectories. Interestingly, we identified a marked decrease in temporal gene expression correlations approximately 60% of the way through both trajectories aligning with visual differences in branch-specific genes and the onset of parenchyma initials bifurcation (Figure S11F, Figure S11G). Further dissection of this time point revealed that a homolog of *ATHB*-*13* (hereafter referred to as *GmATHB13*) was the first TF to be differentially expressed between axis and cotyledon parenchyma at parenchyma initials bifurcation. Interestingly, *ATHB-13* is an HD-Zip I TF previously associated with cotyledon morphogenesis in *Arabidopsis*^72^ and null alleles of *ATHB-13* exhibit increased root length^73^ which is developed from the axis tissue in soybean seed. Thus, we hypothesized that *GmATHB13* acts as a negative regulator of axis development by promoting cotyledon parenchyma identity.

Next, to showcase the power of pseudotime analysis for understanding cellular differentiation, we aimed to characterize the targets and dynamics of GmATHB13 across cotyledon parenchyma development. First, we defined the putative targets of GmATHB13 as the set of expressed genes within the cotyledon parenchyma trajectory with nearby ACRs containing the ATHB13 motif (n=2,177), as well as a set of cotyledon parenchyma expressed control ‘non-target’ genes (n=2,177) lacking ATHB13 motifs within nearby ACRs (Figure 6K). Consistent with the known function of cotyledon parenchyma, expressed genes with accessible ATHB13 motifs were enriched for GO terms related to carbohydrate, polysaccharide, glycogen, and energy reserve metabolic processes (Table S20). We then evaluated expression and TF motif deviation dynamics of ATHB13 in unison with the expression patterns of putative ATHB13 targets and the set of control genes (Figure 6K). *GmATHB13* is initially expressed at low levels and then reaches a peak immediately after the bifurcation point that is followed by a rapid decrease. This indicates that *GmATHB13* expression is tightly correlated with bifurcation of parenchyma initials in a dose-dependent manner. Global chromatin accessibility of the ATHB13 motif increased markedly following the peak of *GmATHB13* expression, suggesting a genome-wide increase in ATHB13 DNA-binding activity that depends on *GmATHB13* transcript levels. Finally, putative ATHB13 targets show higher levels of expression compared to the control set following bifurcation, implicating GmATHB13 as a transcriptional activator. These data suggest that the expression of *GmATHB13* in parenchyma initials above a dosage threshold results in the activation of a gene expression program that promotes cotyledon parenchyma identity.

## Discussion

In-depth knowledge of cell-type resolved transcriptional regulatory programs is essential for gene function studies and gene regulatory network discovery, which are key to both developmental biology and crop improvement.^74^ Here, we constructed a comprehensive single-cell CRE and gene expression atlas by integrating single-cell genomic and spatial technology, profiling 316,358 cells across ten primary tissues in soybean. We assessed the accessibility of approximately 300,000 ACRs across 103 cell types, measuring the cell-type-specific CRE activity that drives dynamic gene expression from the soybean genome. This ACR atlas represents a valuable resource for the soybean community to understand the molecular patterns underlying cell-type diversification in soybean. Additionally, this work provides a framework for constructing cell-type-specific *cis*-regulatory maps for other non-model species lacking known functional marker genes.

The identification of CREs and TFs with cell-type-specific activities provided a comprehensive roadmap for studying regulatory dynamics across cell types and developmental stages. Our results integrate single-cell chromatin and transcriptome data and allowed us to find soybean gene regulatory networks that recapitulate those identified in other species. Notably, using data from infected cells in developing nodules, we successfully *de novo* identified four TF motifs of known master regulators of nodulation and identified their TF binding sites in the ACRs of their target genes (Figure 4H-K). We also discovered two novel infected cell-specific TF motifs that underpin unknown roles in symbiotic nitrogen fixation that can be explored to find novel TFs needed for nitrogen fixation. These results demonstrate how integrating single-cell transcriptome and chromatin accessibility data can discover new cell type regulators and their gene regulatory networks.

The endosperm is fundamental to soybean seed development, providing nutrition through the high expression of nutrition transporter genes.^54^ Among the 80 sucrose transporters in the soybean genome the endosperm expressed *GmSWEET15a* and *GmSWEET15b* played significant roles in increasing seed size and oil content in soybean domestication and modern breeding^29^. However, their precise expression patterns across seed development were unclear, as was whether other transporters have similar expression patterns that could be exploited in soybean breeding. Our results suggest that *GmSWEET15a* and *GmSWEET15b* are specifically expressed in the peripheral endosperm and are upregulated during seed development. Along with these two genes, a group of 13 sucrose transporters showed similar expression patterns and shared the same motif binding site of DOF transcription factors in their candidate CREs. These DOF regulated late stage of peripheral endosperm sugar transporters may likewise affect seed size and oil content (Figure 5P). Interestingly, OsDof11 also controls six sugar transporter genes by directly binding to their promoters and regulating rice seed size^75^, suggesting that the DOF-SWEET gene regulation may be conserved across monocots and dicots^75^ These findings highlight the value of our dataset for precisely studying gene function and positioning genes within transcriptional regulatory networks.

The seed is the agronomic product of soybean, and despite significant efforts studying soybean seeds,^28,76,77^ the gene regulatory networks underpinning seed development are not well characterized. By producing single-cell transcriptome and chromatin accessibility data across seed development, we provide the resources needed to identify these seed developmental regulatory networks. Exemplifying this, we identified the main embryo cell lineages and constructed a comprehensive pseudotime trajectory for embryogenesis, successfully finding known transcriptional regulators, such as *PDF1* and *MP.*^70^ A detailed comparison of regulatory dynamics between cotyledon and axis parenchyma lineages revealed that differential expression of *GmATHB-13* coincides with the lineage bifurcation between axis and cotyledon parenchyma. *Arabidopsis ATHB-13* regulates cotyledon morphogenesis, and genes containing *ATHB-13* motifs are enriched in carbohydrate and polysaccharide metabolism and biosynthesis, matching the expected functions of cotyledon parenchyma cells, which are energy production and nutrition storage. These results suggest that *GmATHB-13* is a good candidate for modifying seed size or composition in soybean, as it may trigger the fate decision between axis and cotyledon parenchyma.

Our analyses are just a starting point, with many other insights to be discovered from these data by exploring the expression patterns and regulatory networks of other genes interest. To facilitate future discovery, we constructed a soybean multi-omic atlas database (https://soybean-atlas.com/), which includes chromatin accessibility and gene expression data for all the cell types explored here. To demonstrate how to explore the gene regulatory network using the database, we created a workflow focusing on predicting the gene regulatory network for LEAFY COTYLEDON1 (LEC1) (Figure S12), a central regulator controlling embryo and endosperm development^78^. We found several interesting observations for *GmLEC1a/b* regulation directly from the database: 1) Two ACRs were identified in the first intron of the paralogs, which were specifically accessible in endosperm and embryo cells; 2) These ACRs captured two motifs consistently enriched in endosperm or embryo cells at three stages of developing seeds: the GmABI3A (ABA INSENSITIVE3a) motif, which controls embryo development and directly binds *GmLEC1*^28^, and the MYB118 motif, which is specifically expressed in endosperm and control endosperm maturation in *Arabidopsis*^47^; 3) *GmABI3a* and its TF motif was mainly expressed and accessible in embryo cells in cotyledon stage seeds, while *GmMYB118a/b* and their TF motif were mainly expressed and accessible in endosperm. Thus, we can propose a model where the specific use of the intronic MYB118 and ABI3 motifs contributes to the expression pattern of *GmLEC1a/b* (Figure S12). The soybean multi-omic atlas is easy to explore via the interactive website, allowing the soybean community to study the gene regulatory networks, at cell-type resolution, for all soybean traits.

Additionally, all preprocessed data matrices, including cell-type-specific ACRs, genes, and TF motifs, are also accessible through The National Center for Biotechnology Information^79^ (NCBI GEO: GSE270392) and SoyBase (https://www.soybase.org/)^80^. We anticipate that the real potential of single-cell methods will extend beyond aiding gene function studies and uncovering regulatory networks - It will involve combining single-cell gene regulatory atlases with machine learning and high-throughput perturbation techniques, to achieve a profound and predictive understanding of gene regulation throughout plant development to improve crop performance.

## Acknowledgement

We would like to acknowledge Dr. Jianxin Ma for sharing the *Bradyrhizobium japonicum* strain USDA110, Dr. Hang Yin for providing access to their cryostat, Dr. Aaron Mitchell for access to their microscope. We especially express our appreciation to Dr. Robert B. Goldberg, Dr. John J. Harada and Dr. Matteo Pellegrini for their pioneering work in “GENE NETWORKS IN SEED DEVELOPMENT” (http://seedgenenetwork.net/), which was critical for evaluating the quality of the single-cell genomic data and analysis. This research was supported by the United Soybean Board (2432-201-0102) and the National Science Foundation (IOS-1856627) to RJS and the National Institute for General Medical Sciences of the National Institutes of Health (R00GM144742) to A.P.M.

## Author contributions

R.J.S. and X.Z. designed the research. X.Z. and Z.L. performed the experiments. X.Z., Z. L., A.P.M., H.Y., H.J., S.B., J.P.M., M.A.M. and R.J.S. analyzed the data. X.Z., Z.L., A.P.M., and R.J.S. wrote the manuscript. The authors read and approved the final manuscript.

## Supplemental information

Table S1. Summary of quality control metrics for the scATAC-seq libraries, related to Figure 1, Figure S1.

Table S2. Summary of quality control metrics for the snRNA-seq libraries, related to Figure 1, Figure S1.

Table S3. Sequencing and barcode meta data for scATAC-seq, related to Figure 1, Figure S1.

Table S4. Sequencing and barcode meta data for snRNA-seq, related to Figure 1, Figure S1.

Table S5. Literature-derived marker genes and cell type metrics, related to Figure 1, Figure S3.

Table S6. Summary of quality control metrics for the spatial transcriptome libraries, related to Figure 2.

Table S7. The *de novo* cell-type specific marker genes identified from the snRNA-seq data, related to Figure 2.

Table S8. The *de novo* cell-type specific marker genes identified from the spRNA-seq data, related to Figure 2, Figure S10.

Table S9. Significant association between ACR and flanking genes, related to Figure 3, Figure S8.

Table S10. Enriched motif list for activating ACRs and repressing ACRs, related to Figure 3, Figure S8.

Table S11. List for cell-type-specific ACR across all cell types, related to Figure 4, Figure S9.

Table S12. Enriched motif for cell-type-specific ACRs, related to Figure 4, Figure S9.

Table S13. *de novo* motif enrichment for infected cell-specific ACRs, related to Figure 4, Figure S9.

Table S14. GO enrichment for three sub-cell types of endosperm at the cotyledon stage, related to Figure 5.

Table S15. Sequencing and barcode meta data for integrated endosperm cells from scATAC-seq and snRNA-seq, related to Figure 5, Figure S10.

Table S16. List of pseudotime genes for peripheral endosperm, related to Figure 5.

Table S17. List of pseudotime genes for micropylar endosperm, related to Figure 5.

Table S18. GO enrichment list for pseudotime genes of micropylar and peripheral, related to Figure 5.

Table S19. GO enrichment list for ATHB13 binding gene in embryo, related to Figure 6.

## Experimental model and subject details

### Growth conditions

The soybean seeds of the Williams 82 genotype were obtained from the USDA National Plant Germplasm System (https://npgsweb.ars-grin.gov) and sown in Sungro Horticulture professional growing mix (Sungro Horticulture Canada Ltd.). For libraries derived from leaf, hypocotyl, nodule, and seed-related tissues, the plants were grown in a greenhouse under a 50/50 mixture of 4100K (Sylvania Supersaver Cool White Deluxe F34CWX/SS, 34W) and 3000K (GE Ecolux with Starcoat, F40CX30ECO, 40W) lighting, with a photoperiod of 16 hours of light and 8 hours of dark. The temperature was maintained at approximately 25°C during light hours, with a relative humidity of approximately 54%.

#### Soybean leaves

For each sample, approximately 6 leaves with a 1 cm diameter were harvested between 8 and 9 AM, ten days after sowing. Fresh tissue was used to construct bulk ATAC-seq, scATAC-seq and snRNA-seq libraries.

#### Soybean hypocotyls

For each sample, approximately 4 hypocotyls were harvested between 8 and 9 AM, seven days after sowing. Fresh tissue was used to construct scATAC-seq and snRNA-seq libraries.

#### Soybean roots

Soybean root samples were obtained as follows: soybean seeds were sterilized with 70% ethanol for 1 minute. After removing the ethanol solution, the seeds were treated with 10% bleach for 5 minutes, followed by five washes with autoclaved Milli-Q water. The sterilized seeds were then sown on mesh plates with half-strength MS media (Phytotech Laboratories, catalog: M519) and wrapped in Millipore tape. Plates were incubated in a Percival growth chamber with a photoperiod of 16 hours of light and 8 hours of dark. The growth chamber temperature was set to 25°C with a relative humidity of approximately 60%. For each sample, approximately 5 whole roots were harvested between 8 and 9 AM, seven days after sowing. Fresh tissue was used to construct scATAC-seq and snRNA-seq libraries.

#### Soybean nodules

Soybean nodules were induced following a previously described soil-free method for producing root nodules in soybean.^81^ Briefly, seeds were germinated in sterilized germination paper (Anchor Paper Company, St Paul, MN, USA) wetted with autoclaved water for 10 days. The roots were then infected with *Bradyrhizobium japonicum* strain USDA110 to produce nodules. Roots with nodules approximately 1 mm in diameter were collected 15 days post-inoculation (dpi), and root tips were removed (Figure 4F). The tissue was flash-frozen in liquid nitrogen and stored at -80°C. For each sample, approximately 10 tissues were used for scATAC-seq and snRNA-seq preparation.

#### Soybean pods

For each sample, approximately 20 whole pods, each 5 mm in length, were harvested between 8 and 9 AM in the greenhouse. Fresh tissue was used to construct scATAC-seq and snRNA-seq libraries.

#### Soybean seeds

Seed stages were determined according to previously described methods and standards.^82^ Specifically, seed lengths for the globular, heart, cotyledon, and early maturation stages were 1.0 mm, 2 mm, 3 mm, and 7 mm, respectively. Seeds at the middle maturation stage weighed about 200-250 mg. Fresh tissue was used to construct scATAC-seq and snRNA-seq libraries for all seed tissues.

### scATAC-seq library preparation

Nuclei isolation and purification were performed as described previously.^51^ Briefly, the tissue was finely chopped on ice for approximately 2 minutes using 600 μL of pre-chilled Nuclei Isolation Buffer (NIB: 10 mM MES-KOH at pH 5.4, 10 mM NaCl, 250 mM sucrose, 0.1 mM spermine, 0.5 mM spermidine, 1 mM DTT, 1% BSA, and 0.5% Triton X-100). After chopping, the mixture was passed through a 40-μm cell strainer and centrifuged at 500 rcf for 5 minutes at 4°C. The supernatant was carefully decanted, and the pellet was reconstituted in 500 μL of NIB wash buffer (10 mM MES-KOH at pH 5.4, 10 mM NaCl, 250 mM sucrose, 0.1 mM spermine, 0.5 mM spermidine, 1 mM DTT, and 1% BSA). The sample was filtered through a 10-μm cell strainer and gently layered onto 1 mL of 35% Percoll buffer (35% Percoll mixed with 65% NIB wash buffer) in a 1.5-mL centrifuge tube. The nuclei were centrifuged at 500 rcf for 10 minutes at 4°C. After centrifugation, the supernatant was carefully removed, and the pellets were resuspended in 10 μL of diluted nuclei buffer (DNB, 10X Genomics Cat# 2000207). Approximately 5 μL of nuclei were diluted tenfold, stained with DAPI (Sigma Cat. D9542), and the nuclei quality and density were evaluated using a hemocytometer under a microscope. The original nuclei were then diluted with DNB buffer to a final concentration of 3,200 nuclei per μL. Finally, 5 μL of nuclei (16,000 nuclei in total) were used as input for scATAC-seq library preparation.

scATAC-seq libraries were prepared using the Chromium scATAC v1.1 (Next GEM) kit from 10X Genomics (Cat# 1000175), following the manufacturer’s instructions (10x Genomics, CG000209_Chromium_NextGEM_SingleCell_ATAC_ReagentKits_v1.1_UserGuide_RevE). Libraries were sequenced on an Illumina NovaSeq 6000 in dual-index mode with eight and 16 cycles for i7 and i5 indexes, respectively.

### Bulk ATAC-seq library preparation

Nuclei isolation followed the exact procedure used for scATAC-seq, and the library preparation strictly adhered to the protocol described previously^83^.

### snRNA-seq library preparation

The protocol for nuclei isolation and purification was adapted from a previously described scATAC-seq method. To minimize RNA degradation and leakage, the tissue was finely chopped on ice for approximately 1 minute using 600 μL of pre-chilled Nuclei Isolation Buffer containing 0.4 U/μL RNase inhibitor (Roche, Protector RNase Inhibitor, Cat. RNAINH-RO) and a low detergent concentration of 0.1% NP-40. Following chopping, the mixture was passed through a 40-μm cell strainer and centrifuged at 500 rcf for 5 minutes at 4°C. The supernatant was carefully decanted, and the pellet was reconstituted in 500 μL of NIB wash buffer (10 mM MES-KOH at pH 5.4, 10 mM NaCl, 250 mM sucrose, 0.5% BSA, and 0.2 U/μL RNase inhibitor). The sample was filtered again through a 10-μm cell strainer and gently layered onto 1 mL of 35% Percoll buffer (prepared by mixing 35% Percoll with 65% NIB wash buffer) in a 1.5-mL centrifuge tube. The nuclei were centrifuged at 500 rcf for 10 minutes at 4°C. After centrifugation, the supernatant was carefully removed, and the pellets were resuspended in 50 μL of NIB wash buffer. Approximately 5 μL of nuclei were diluted tenfold and stained with DAPI (Sigma Cat. D9542). The quality and density of the nuclei were evaluated using a hemocytometer under a microscope. The original nuclei were further diluted with DNB buffer to achieve a final concentration of 1,000 nuclei per μL. Ultimately, a total of 16,000 nuclei were used as input for snRNA-seq library preparation.

For scRNA-seq library preparation, we employed the Chromium Next GEM Single Cell 3’GEM Kit v3.1 from 10X Genomics (Cat# PN-1000123), following the manufacturer’s instructions (10xGenomics, CG000315_ChromiumNextGEMSingleCell3-_GeneExpression_v3.1_DualIndex_RevB). The libraries were subsequently sequenced using the Illumina NovaSeq 6000 in dual-index mode with 10 cycles for the i7 and i5 indices, respectively.

### Spatial RNA-seq library preparation

For the spatial RNA-seq experiment, the hypocotyl tissues, the root tissues, and the seed tissues at the heart stage, cotyledon stage, and early maturation stage, matching the stages of the single-cell datasets, were sampled. The tissues were embedded in the Optimal Cutting Temperature (OCT) compound, snap-frozen in a cold 2-methylbutane bath merged in liquid nitrogen, and cryosectioned into 12 um thick slices.

We used the Visium Spatial Gene Expression Kit (10X Genomics, USA) to construct the spatial RNA-seq libraries following the manufacturer’s instructions. The tissue sections were mounted onto the spatial slides, fixed by cold methanol, and stained by 0.05% toluidine blue. The stained tissue sections were imaged using the BZ-X800 fluorescent microscope (Keyence, Japan). To determine the optimal tissue permeabilization time, we performed the Tissue Optimization workflow on a series of digestion times for each tissue type. For the spatial RNA-seq libraries, mRNA was first released according to the optimal permeabilization time, then the spatially barcoded cDNAs were synthesized on the slides. Finally, cDNA were released from the slide and subjected to amplification and library construction, following the manufacturer’s specifications

## Quantification and statistical analysis

### scATAC-seq raw reads processing

The raw data processing followed the previously described method.^19^ In brief, raw BCL files were demultiplexed and converted into fastq format using the default settings of the 10X Genomics tool cellranger-atac makefastq (v1.2.0). Initial read processing, including adaptor/quality trimming, mapping, and barcode attachment/correction, was carried out with cellranger-atac count (v1.2.0) using the soybean William 82 v4 reference genome and the Glycine max organelle genomes (NCBI Reference Sequence: NC_007942.1, NC_020455.1).^84^ Properly paired, uniquely mapped reads with a mapping quality greater than 30 were retained using samtools view (v1.6; -f 3 -q 30) and reads with XA tags were filtered out.^85^ Duplicate fragments were collapsed on a per-nucleus basis using picardtools (http://broadinstitute.github.io/picard/) MarkDuplicates (v2.16; BARCODE_TAG=CB REMOVE_DUPLICATES=TRUE). Reads mapping to mitochondrial and chloroplast genomes were counted for each barcode and then excluded from downstream analysis. Potential artifacts were removed by excluding alignments coinciding with a blacklist of regions exhibiting Tn5 integration bias from Tn5-treated genomic DNA (1-kb windows with greater than 4x coverage over the genome-wide median) and potential collapsed sequences in the reference (1-kb windows with greater than 4x coverage over the genome-wide median using ChIP-seq input). BAM alignments were then converted to single base-pair Tn5 integration sites in BED format by adjusting coordinates of reads mapping to positive and negative strands by +4 and -5, respectively, and retaining only unique Tn5 integration sites for each distinct barcode.

The R package Socrates was used for nuclei identification and quality control.^19^ The BED file containing single base-pair Tn5 integration sites was imported into Socrates along with the soybean GFF gene annotation (Phytozome, version Gmax_508_Wm82.a4.v1) and the genome index file. To identify bulk-scale ACRs in Socrates, the callACRs function was employed with the following parameters: genome size=8.0e8, shift=-75, extsize=150, and FDR=0.1. This step allowed us to estimate the fraction of Tn5 integration sites located within ACRs for each nucleus. Metadata for each nucleus were collected using the buildMetaData function, with a TSS (Transcription Start Site) window size of 1 kb (tss.window=1000). Sparse matrices were then generated with the generateMatrix function, using a window size of 500. High-quality nuclei were identified based on the following criteria: a minimum of 1,000 Tn5 insertion sites per nucleus, at least 20% of Tn5 insertions within 2 kb of TSSs, and at least 20% of Tn5 insertions within ACRs across all datasets. Additionally, a maximum of 20% of Tn5 insertions in organelle genomes was allowed.

For each tissue, integrated clustering analysis of all replicates was performed using the R package Socrates.^19^ For the binary nucleus x window matrix, windows accessible in less than 1% of all nuclei and nuclei with fewer than 100 accessible ACRs were removed using the function cleanData (min.c=100, min.t=0.01). The filtered nucleus x window matrix was normalized with the term-frequency inverse-document-frequency (TF-IDF) algorithm with L2 normalization (doL2=T). The dimensionality of the normalized accessibility scores was reduced using the function reduceDims while removing singular values correlated with nuclei read depth (method=“SVD”, n.pcs=25, cor.max=0.4). The reduced embedding was visualized as a UMAP embedding using projectUMAP (k.near=15). Approximately 5% of potential cell doublets were identified and filtered by performing a modified version of the Socrates workflow on each library separately with the function detectDoublets and filterDoublets (filterRatio=1.0, removeDoublets=T). To address batch effects, we used the R package Harmony with non-default parameters (do_pca=F, vars_use=c(“batch”), tau=5, lambda=0.1, nclust=50, max.iter.cluster=100, max.iter.harmony=50). The dimensionality of the nuclei embedding was further reduced with Uniform Manifold Approximation Projection (UMAP) via the R implementation of projectUMAP (metric=“correlation”, k.near=15). Finally, the nuclei were clustered with the function callClusters (res=0.5, k.near=15, cl.method=3, m.clust=25).

### snRNA-seq raw reads processing

*STARSolo* was used to map the snRNA-seq reads and count the gene features using the soybean genome (William 82 v4).^86^ We specified the following parameters in *STARSolo* to filter the UMI, filter empty cells, and count multi-mapping reads: --soloUMIfiltering MultiGeneUMI_CR, --soloCellFilter EmptyDrops_CR, --soloMultiMappers PropUnique. The filtered expression data was analyzed using the *Seurat* (v4) R package.^26^ Potential low-quality nuclei or empty droplets were filtered. Specifically, cells with gene counts below a threshold calculated as the median gene count minus two times the median absolute deviation, and cells with UMI counts less than the lower 10% percentile of total UMI counts, were filtered out. Additionally, cells with organelle gene counts comprising more than 15% of the total gene count were excluded. The preprocessed datasets were normalized using *SCTransform* before the *RunPCA* for principal component analysis (PCA). Subsequently, the doublets were identified by the *DoubletFinder* R package, and removed from downstream analysis. We prepared two replicates for each library and integrated them using the *Harmony* R package.^87^ The integrated dataset was then processed using *RunUMAP* (reduction = “harmony”, dims = 1:20) for Uniform Manifold Approximation and Projection (UMAP) dimension reduction, *FindNeighbors* (reduction = “harmony”, dims = 1:30) to obtain the Nearest-neighbor graph, and *FindClusters* to identify distinct cell populations. Different resolutions were selected to classify cell types in varying tissue types. We used *FindSubCluster* to identify the sub-clusters according to the specificity of marker genes.

### spRNA-seq reads processing

We used Space Ranger (10X Genomics) to map the spRNA-seq reads to the soybean genome and to count gene expression. The filtered gene expression matrix was analyzed using the *Seurat* (v4) R package.^26^ All the datasets were analyzed using *SCTransform* and *RunPCA*. To remove the batch effect for replicates placed in different spatial capture areas, we used the *Harmony* R package to integrate the replicates and analyzed it using *RunUMAP* (reduction = “harmony”, dims = 1:20) and *FindNeighbors* (reduction = “harmony”, dims = 1:20). We used *FindClusters* to identify cell clusters and *FindSubCluster* to identify the subclusters for specific cell types. Various resolutions were used to identify the cell clusters in distinct types of tissues.

### Integration of snRNA-seq and spRNA-seq

We applied the ‘anchor’-based integration method from *Seurat* to integrate the snRNA-seq and spRNA-seq datasets.^88^ First, we used *FindTransferAnchors* (normalization.method=“SCT”) to find the anchors between the reference dataset (snRNA-seq) and the query dataset (spRNA-seq). These anchors were used to calculate the prediction scores of each snRNA-seq cell type for the spRNA-seq using the *TransferData* (dims = 1:30).

### *De novo* marker identification

After cell type annotation, we identified the *de novo* marker genes using the *FindAllMarkers* (test.use=”wilcox”, logfc.threshold = 1, only pos=T, min.pct = 0.1) from the Seurat R package. Then we took the top 50 most up-regulated genes and filtered them by adjusted p-value>0.00001 and log2FC>2 to obtain the significant marker genes.

### Cell-type annotation for snRNA-seq

To assign cell types to each cluster, we used a combination of marker gene-based annotation and gene set enrichment analysis. Initially, we compiled a list of known cell-type-specific marker genes known to localize to discrete cell types or domains expected in the sampled tissues based on an extensive review of the literature (Table S5). And the ortholog list for *Arabidopsis* and soybean was downloaded from PANTHER (v18.0)^89^. Gene expression was calculated using the UMI counts in the gene body and aggregating all nuclei in a cluster, then the raw counts matrix was normalized with the CPM function in edgeR. The *Z*-score was calculated for each marker gene across all cell types using the scale function in R, and key cell types were assigned based on the most enriched marker genes with the highest *Z*-score. Ambiguous clusters displaying similar patterns to key cell types were assigned to the same cell type as the key cell types, reflecting potential variations in cell states within a cell type (Figure S3). To aid visualization, we smoothed normalized gene accessibility scores by estimating a diffusion nearest neighbor graph.^19^

For soybean seed tissue, the cpm normalized matrix was also mapped to the subregion by checking the correlation with the laser capture microdissection (LCM) RNA-seq dataset (http://seedgenenetwork.net/seeds). With this approach, we could clearly identify the seed coat, endosperm, and embryo regions, which confirmed our cell type annotation. There were no available markers for seed coat endothelium and seed coat inner integument, so these two cell types were annotated based on specific high correlations with the LCM dataset (Figure S2 E,F).

For gene set enrichment analysis, we used the R package fgsea, following a methodology described previously.^19,90^ Firstly, we constructed a reference panel by uniformly sampling nuclei from each cluster, with the total number of reference nuclei set to the average number of nuclei per cluster. Subsequently, we aggregated the UMI counts across nuclei in each cluster for each gene and identified the differential expression profiles for all genes between each cluster and the reference panel using the R package edgeR.^91^ For each cluster, we generated a gene list sorted in decreasing order of the log2 fold-change value compared to the reference panel and utilized this list for gene set enrichment analysis. We excluded GO terms with gene sets comprising less than 10 or greater than 600 genes from the analysis, and GO terms were considered significantly enriched at an FDR < 0.05 with 10,000 permutations. The cell type annotation was additionally validated by identifying the top enriched GO terms that align with the expected cell type functions.

### Cell-type annotation for scATAC-seq

A similar approach used for snRNA-seq cell type annotation was applied to scATAC-seq with minor optimizations. Specifically, the gene chromatin accessibility score, rather than gene expression, was calculated using the Tn5 integration number in the gene body, a 500 bp upstream region, and a 100 bp downstream region. The raw counts were then normalized with the cpm function in edgeR. Cell types were assigned to each cluster following the snRNA-seq annotation process, including evaluating marker gene performance and GO enrichment profiles.

For tissues with both snRNA-seq and scATAC-seq data, we further confirmed the cell annotations by integrating the two modalities using the *Seurat* workflow (v4.0.4).^26^ Briefly, the gene chromatin accessibility score was normalized and scaled with the functions NormalizeData and ScaleData. The function FindTransferAnchors was used for canonical correlation analysis (CCA) to compare the scATAC-seq gene score matrix with the scRNA-seq gene expression matrix and to find mutual nearest neighbors in low-dimensional space. Annotations from the scRNA-seq dataset were then transferred onto the scATAC-seq cells using the TransferData function, and prediction scores less than 0.5 were filtered out. This approach allowed us to match and validate cell types across the two modalities, and we observed a median prediction score of 0.75 across the seven tissues (Figure S2G-I). Finally, we calculated the Pearson correlation coefficient with the top 1,000 variable genes from snRNA-seq, which ranged from 0.4 to 0.7 for the same cell type across the two modalities, similar to observations from other studies (Figure S4).^19,68,92^

### ACR identification

Following cell clustering and annotation, peaks were identified using all Tn5 integration sites for each cluster by running MACS2 with non-default parameters: --extsize 150 --shift -75 --nomodel --keep-dup all.^93^ To account for potential bias introduced by read depth, we adjusted the q-value cutoffs based on the total Tn5 integration number in each cell type as follows: for less than 10 million integrations, we used --qvalue 0.1; for 10-25 million, we used 0.05; for 25-50 million, we used 0.025; for 50-100 million, we used 0.01; and for more than 100 million, we used 0.001. Peaks were then redefined as 500-bp windows centered on the peak coverage summit. To consolidate information across all clusters, we concatenated all peaks into a unified master list using a custom script.^19^ The peak chromatin accessibility score was calculated based on the Tn5 integration count within the peak and then normalized using the cpm function in edgeR.^91^ ACRs with less than 4 CPM in all cell types were removed from downstream analysis. We also used the same method described above to identify the ACRs for bulk ATAC-seq data.

### Predicting the functions of ACRs

We hypothesized that the ACRs only control the flanking genes and used a correlation-based approach to predict the function of the ACRs. Firstly, we created the count matrix of the ACRs and gene expression across 66 main shared cell types between scATAC-seq and snRNA-seq. The count matrix was then normalized using the cpm function in edgeR and the normalize.quantiles function in preprocessCore (v1.57.1).^94^ For each test, we calculated the Spearman correlation between the ACRs accessibility and gene expression, shuffling the ACRs accessibility and gene expression 1,000 times to obtain a p-value for each correlation. This allowed us to compute the p-value for each correlation and adjust for multiple hypotheses using the Benjamini-Hochberg procedure (FDR). We then selected all correlations below -0.25 and above 0.25 with an FDR below 0.05. To simplify the ACRs function, we hypothesized that one ACR controls one gene. For ACRs associated with multiple genes, we filtered the associations based on the following criteria: (i) Kept the best association with the highest correlation if all the associations were genic and proximal. (ii) Kept the best association with the highest correlation if all the associations were distal. (iii) If the associations were a mix of distal or genic and proximal, we only kept distal associations with higher correlation than the genic or proximal associations. Finally, the ACRs with all positive correlations with a flanking gene were predicted as activating ACRs, and the ACRs with all negative correlations with a flanking gene were predicted as repressing ACRs. About 3.9% of ACRs had both negative and positive correlations with a flanking gene, and these ACRs with ambiguous functions were removed from downstream analysis.

### Identification of cell-type-specific ACRs

To identify the cell-type-specific ACRs, we first identified the differentially accessible chromatin regions for each cell type in the tissue. Specifically, for each cell type, we constructed a reference panel by uniformly sampling nuclei from other cell types, with the total number of reference nuclei set to the number of nuclei in the tested cell type. Subsequently, we aggregated the Tn5 integration counts across nuclei in the cell type and identified the differential accessibility profiles for all ACRs between each cell type and their reference panel using the R package edgeR. High accessible ACRs in a cell type with a fold change > 4 and p-value < 0.05 were aggregated in the tissue. ACRs identified as highly accessible in at most two cell types were retained as cell-type-specific ACRs in the tissue.

### TF Motif deviations score calculation

TF motif deviation scores of specific TF motifs among nuclei were estimated using chromVAR (Schep et al., 2017) with the non-redundant core plant PWM database from JASPAR2022.^95^ The input matrix for chromVAR was filtered to retain ACRs with a minimum of 10 fragments and cells with at least 100 accessible ACRs. We applied smoothing to the bias-corrected motif deviations for each nucleus, integrating them into UMAP embedding for visualization, like the method used for visualizing gene body chromatin accessibility.

### Motif enrichment

Firstly, TF motif occurrences in all ACRs were identified with fimo from the MEME suite toolset (ref) using position weight matrices (PWM) from the non-redundant core plant motif database in JASPAR 2024.^45,96^ To test the motif enrichment in the cell-type-specific ACRs, we compared the motif distribution in the ctACRs and a control set of “constitutive” ACRs, which varied the least and were broadly accessible across cell types (fold change < 2 and p-value > 0.1), using Fisher’s exact test (alternative = ‘greater’) for each cell type and motif. To control for multiple testing, we used the Benjamini-Hochberg method to estimate the FDR, considering tests with FDR < 0.05 as significantly different between the cell-type-specific ACRs and constitutively accessible regions. To test the motif enrichment in the activating ACRs and repressing ACRs, we compared the motif distribution in the activating ACRs and repressing ACRs using Fisher’s exact test (alternative = ‘greater’) for each motif. Motifs with a p-value less than 0.01 were considered significantly enriched.

### *De novo* TF motif enrichment

To identify novel motifs in the cell-type-specific ACRs, we first created a control set by randomly selecting the same number of cell-type-specific ACRs from the “constitutive” ACRs described above, ensuring that they had a similar GC content ratio to the test set. De novo motif searches in cell-type-specific ACRs were performed using XSTREME version 5.5.3 within the MEME suite package (v5.5.0) with the non-default parameter “--maxw 30,” and we provided the known motifs from the non-redundant core plant motif database in JASPAR 2024 or collected from the literature.^97^

### Embryo scATAC-seq and scRNA-seq clustering

To chart the dynamics of chromatin accessibility and transcription during embryogenesis, we first collected all scATAC-seq and snRNA-seq nuclei with embryo cell type annotations from the four matched seed developmental time points (Globular, Heart, Cotyledon, and Early Maturation stages), and re-clustered scATAC-seq and snRNA-seq nuclei, independently.

For the snRNA-seq data set, we first partitioned the nuclei x gene matrix corresponding specifically to embryo cell types and removed genes expressed in less than 0.1% of nuclei. To remove outlier nuclei, we then selected nuclei with at least 100 unique expressed genes and less than 10,000 unique expressed genes. The sparse gene x nuclei matrix was then processed with the R package, *Seurat* (v5.0.1) by first log-normalizing counts using *NormalizeData* with default parameters.^98^ We scaled the normalized counts with *ScaleData* and regressed out effects from variation in the log-scaled UMI counts and percent UMIs mapping to organeller genes. The scaled matrix was then used to identify variable features via *FindVariableFeatures* with non-default parameters (selection.method=”mean.var.plot”, dispersion.cutoff=c(0.5, Inf), mean.cutoff=c(0.0125,3)). To reduce the dimensionality of the nuclei x gene matrix, we ran principal component analysis with *RunPCA* to identify the top 20 PCs. The reduced embedding was used as input for UMAP from the *uwot* R package (min_dist=0.01, n_neighbors=30, metric=”cosine”). We then generated a neighborhood graph with *FindNeighbors* with non-default parameters (dims=1:20, nn.esp=0, k.param=30, annoy.metric=”cosine”, n.trees=100, prune.SNN=1/30, l2.norm=T). Finally, we identified clusters using the *FindClusters* function with resolution=1 and the leiden algorithm (algorithm=4). Cluster cell types were derived from the prior annotation strategy and validated using marker gene expression profiles from the new clustering results (Table S5).

To recluster the scATAC-seq embryo nuclei, we first partitioned the nuclei x ACR matrix specifically for nuclei labeled as embryo cell types from the prior annotation. All downstream scATAC-seq analyses were conducted inside the *Socrates* framework unless otherwise noted. Nuclei with less than 100 unique accessible chromatin regions were removed and ACRs that were accessible in less than 1% of nuclei were also excluded using the function *cleanData* (min.c=100, min.t=0.01). The nuclei x ACR matrix was normalized by TFIDF followed by taking the L2 norm of each nucleus with the function *tfidf* and non-default parameters (doL2=T). To reduce the dimensionality of this matrix, we performed Singular Value Decomposition (SVD), taking the top 25 singular values after removing singular values correlated with per-nucleus read depths greater than 0.5, and L2 normalizing the components via non-default parameters of the function *reduceDims* (n.pcs=25, method=”SVD”, cor.max=0.5, scaleVar=T, doL2=T). The reduced matrix was then projected into two-dimensions with *projectUMAP* with non-default settings (metric=”cosine”, k.near=15). To identify clusters, we generated a shared neighborhood graph and clustered the data using leiden with the function *callClusters* with non-default parameters (res=0.5, k.near=15, cleanCluster=T, cl.method=4, e.thresh=3, m.clust=25, min.reads=5e5) to remove UMAP outliers and clusters with less than 25 nuclei and a total read depth of 500,000. Cell type annotations for each cluster were determined similarly as for the snRNA-seq clustering results.

### Embryo scATAC-seq and snRNA-seq integration

To integrate the scATAC-seq and snRNA-seq nuclei, we first partitioned three matrices (nuclei x gene accessibility, nuclei x ACR, and nuclei x gene expression) to specifically retain embryo nuclei from the scATAC-seq and snRNA-seq clustering results from above. The integration was performed using the unshared features iNMF workflow from the R package, *liger*.^99^ Specifically, we normalized the nuclei x ACR matrix by *tfidf* (*Socrates*) followed by the *normalize* function of *liger* with default settings. The normalized nuclei x ACR slot was then rescaled such that the sum of all accessible regions for a given barcode was 1. Using the *Seurat* framework, we then identified the top 2,000 most variable features using *FindVariableFeatures* with non-default parameters (selection.method=”vst”, nfeatures=2000). The normalized nuclei x ACR matrix was scaled using *scaleNotCenter* and stored as the set of unshared features for downstream integration.

Focusing on the matrices with the shared feature set (geneIDs) between scATAC-seq and snRNA-seq, we selected genes from each modality within the inner 98% quantile of each distribution and retained the intersected genes. The nuclei x gene activity and nuclei x gene expression matrices were normalized using the default settings of the *normalize* function. Variable genes were selected using *selectGenes* with var.thresh=0.1, datasets.use=”RNA”, unshared=TRUE, unshared.datasets=list(2), unshared.thresh=0.2 parameters. The normalized matrices were scaled with *scaleNotCenter* with default settings. The integration was performed with the function *optimizeALS* by setting k=30, use.unshared=TRUE, max_iters=30, and thresh=1e-10. Finally, the integrated embedding was quantile normalized with the function *quantile_norm* setting the reference data set to the snRNA-seq modality.

Using the integrated embedding based on the snRNA-seq nuclei as a reference, we then aimed to impute scATAC-seq modalities on to the snRNA-seq nuclei. To accomplish this, we ran the function *imputeKNN* from the *liger* package to impute motif deviation scores and ACR normalized chromatin accessibility values from the scATAC-seq nuclei onto the snRNA-seq nuclei using default parameters. This results in estimates of gene expression, chromatin accessibility, and motif deviation scores for an individual snRNA-seq barcode.

### Inferred developmental age of embryo nuclei

The time-series nature of the four seed developmental stages of our data lends itself to precise inference of developmental age using model-based approaches.^68^ To simplify interpretation, we focused on the snRNA-seq embryo nuclei across the four developmental stages. Starting from the raw nuclei x gene counts matrix, we log-transformed counts and scaled the resulting values such that the sum across all genes was equal to 10,000 for each barcode. We then downsampled each stage to have the same number of nuclei. Using the R package, *caret*, we partitioned the downsampled nuclei into training and test sets via the function *createDataPartition* with non-default parameters (seed_stage, p=10/11, list=F, times=1). We then trained a linear regression model with a LASSO penalty and 10-fold cross-validation using the *cv.glmnet* function from the R package, *glmnet,* on gene expression profiles for seed stage. The model was then used to collect gene coefficients and continuous developmental age predictions from the entire data set.

### Trajectory analysis

Pseudotime trajectory analysis for each trajectory outlined in Figure 5 H,I and Figure 6E was performed similar to a previously published approach.^19^ Specifically, we ran the function *calcPseudo* with cell.dist1=0.95 and cell.dist2=0.95 from the github repository (https://github.com/plantformatics/maize_single_cell_cis_regulatory_atlas), resulting in pseudotime estimates for individual nuclei for a specific developmental branch. We then identified genes with significant gene expression variation across each trajectory using the function *sigPseudo2* from the same github repo. For visualization, gene expression scores across pseudotime for significantly variable genes were smoothed using predictions on 500 equally spaced bins from a generalized additive model as previously shown.^19^

To identify TFs associated with gene expression variation across pseudotime during Cotyledon parenchyma development, we performed a Pearson’s correlation analysis between TF motif deviations and genes with significant pseudotime variance. TF modules were clustered using k-means, where the final k=8 was selected based on the elbow and silhouette approaches.

## Data and code availability

All datasets generated in this study have been deposited at GEO (Accession number: GSE270392) and are publicly available as of the date of publication.

All original code has been deposited at Github (https://github.com/schmitzlab/soybean_atlas).

Any additional information required to reanalyze the data reported in this paper is available from the lead contact upon request.

## Supplementary

**Figure S1.**
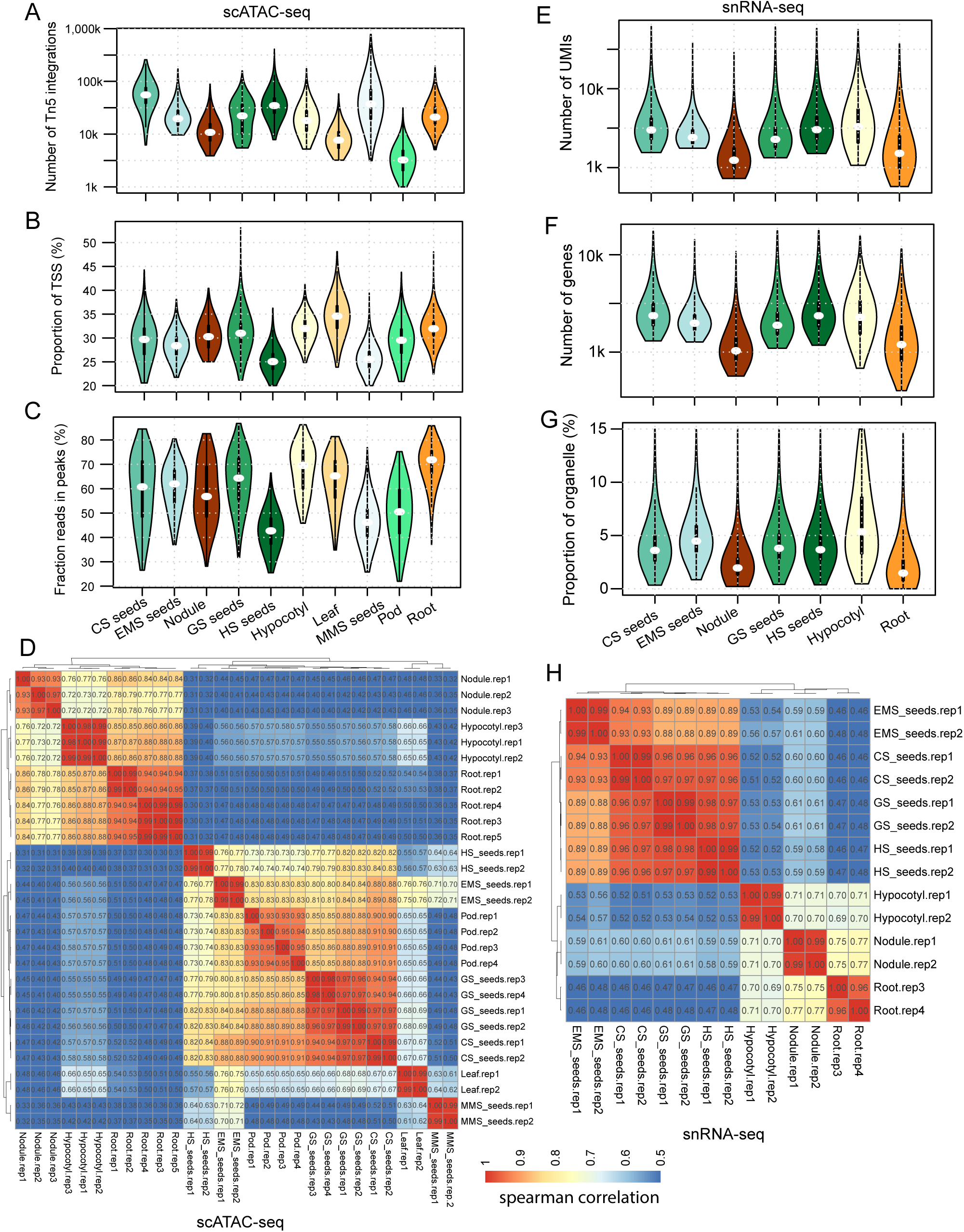
Evaluation and quality control of soybean scATAC-seq and snRNA-seq, related to Figure 1. **(A-D)** Quality control of scATAC-seq: Distribution of unique Tn5 integration sites per nucleus across ten tissues (A); Distributions of the proportion of Tn5 integration sites within the promoter regions, encompassing the 1-kb flanking regions around gene transcription start sites (TSSs) (B); Distributions of the proportion of Tn5 integration sites within peaks per nucleus (C); Spearman correlation coefficient heatmap among all scATAC-seq libraries (D). **(E-H)** Quality control of snRNA-seq: Distribution of total number of UMI (D); Distribution of number of detected genes (E); Distribution of the proportion of reads from organelle (F); Spearman correlation coefficient heatmap among all snRNA-seq libraries (H).

**Figure S2.**
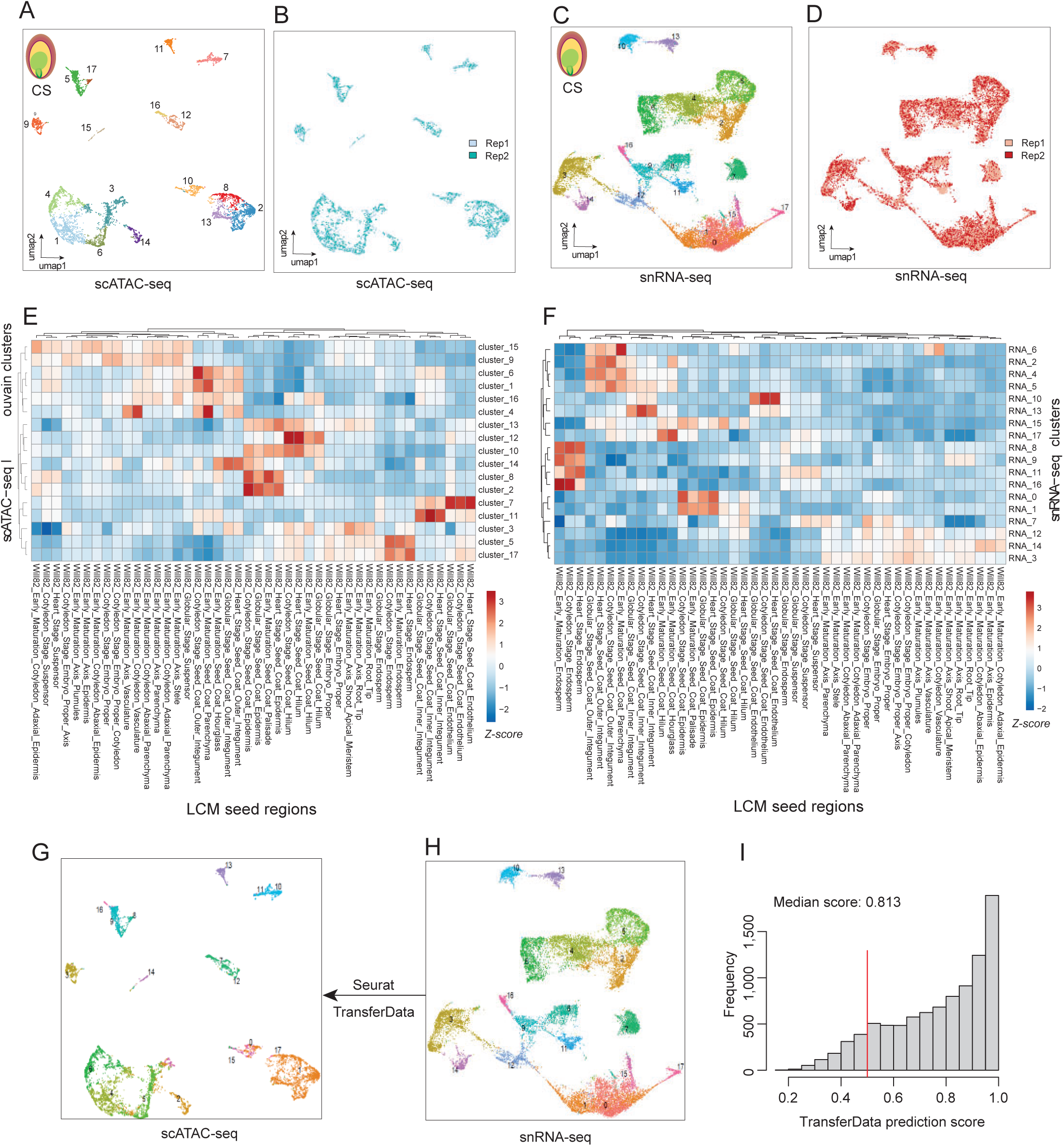
Cell type clustering and initial annotation for soybean seeds at cotyledon stage, related to Figure 1. (A-B) UMAP embeddings for scATAC-seq overlaid with cluster id (A) or library replicates (B). (C-D) UMAP embeddings for snRNA-seq overlaid with cluster id (C) or library replicates (D). (E) Z-score heatmap of spearman correlation coefficient across all laser capture microdissection (LCM) RNA-seq datasets and scATAC-seq clusters. (F) Z-score heatmap of spearman correlation coefficient across LCM RNA-seq datasets and snRNA-seq clusters. (G) UMAP embeddings for scATAC-seq (G) overlaid predicted cluster id in snRNA-seq. (H) UMAP embeddings for snRNA-seq overlaid with raw cluster id. (I) Frequency distribution of max prediction score of snATAC-seq nuclei from the TransferData function in *Seurat*.

**Figure S3.**
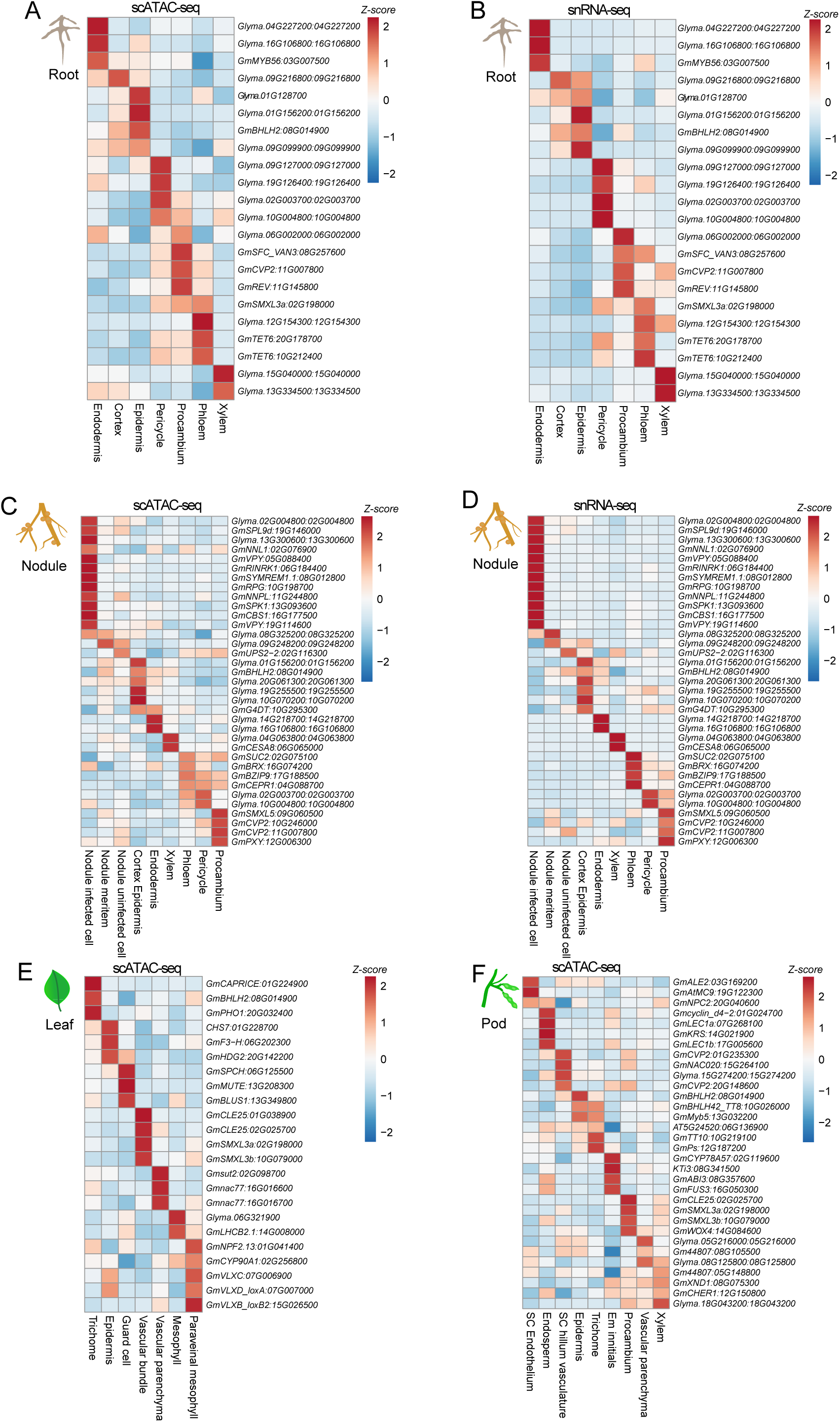
Marker-based annotation for scATAC-seq and snRNA-seq, related to Figure 1. (A-B) *Z*-score heatmap of gene accessibility (A) and gene expression (B) for representative marker genes across shared cell types in soybean roots. (C-D) *Z*-score heatmap of gene accessibility (C) and gene expression (D) for representative marker genes across shared cell types in soybean nodules. (E-F) *Z*-score heatmap of gene accessibility for representative marker genes across cell types in soybean leaves (E) and pods (F).

**Figure S4.**
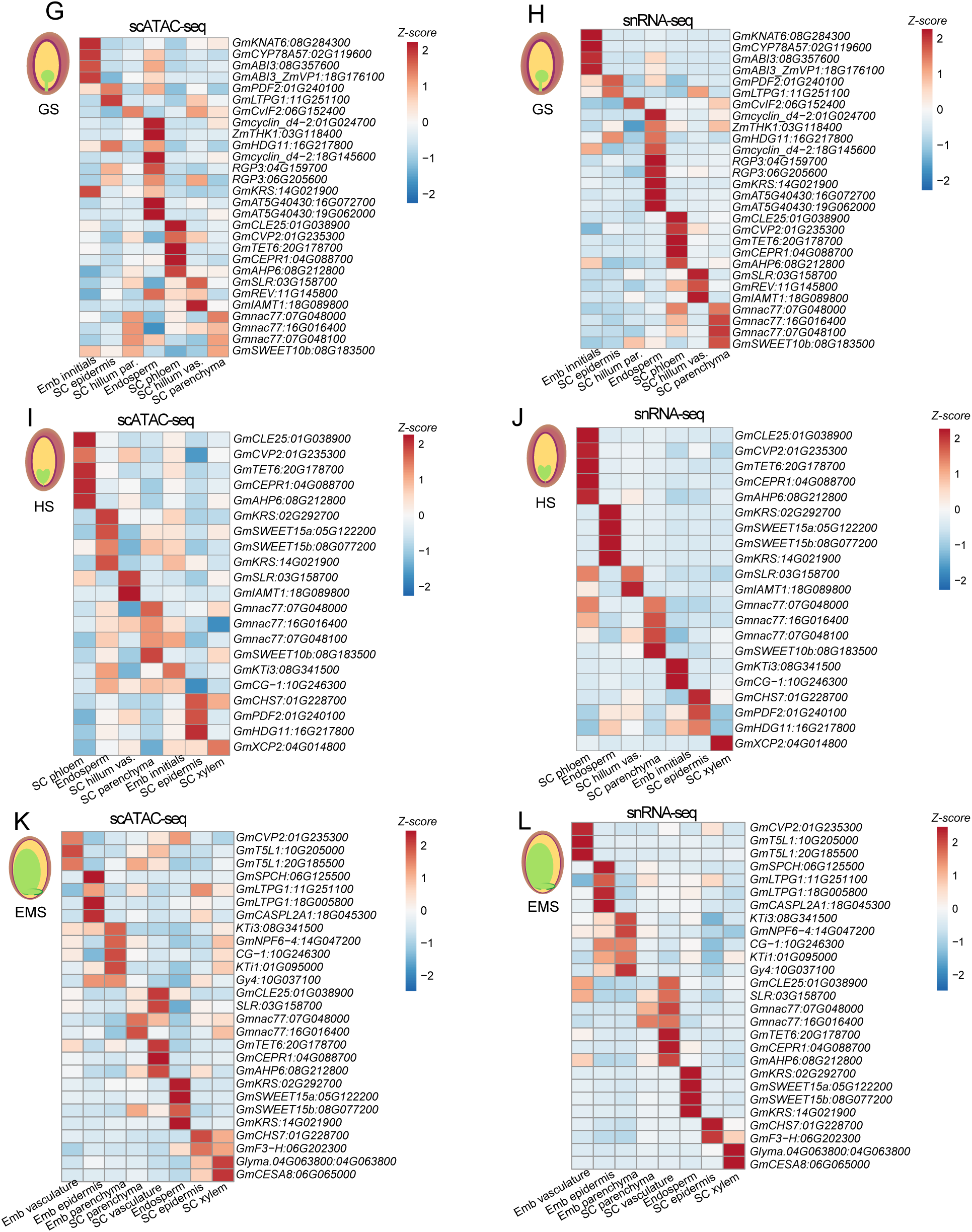
Marker-based annotation for scATAC-seq and snRNA-seq, related to Figure 1. (G-H) *Z*-score heatmap of gene accessibility (G) and gene expression (H) for representative marker genes across shared cell types in soybean seeds at globular stage. (I-J) *Z*-score heatmap of gene accessibility (I) and gene expression (J) for representative marker genes across shared cell types in soybean seeds at heart stage. (K-L) *Z*-score heatmap of gene accessibility (K) and gene expression (L) for representative marker genes across shared cell types in soybean seeds at early maturation stage.

**Figure S5.**
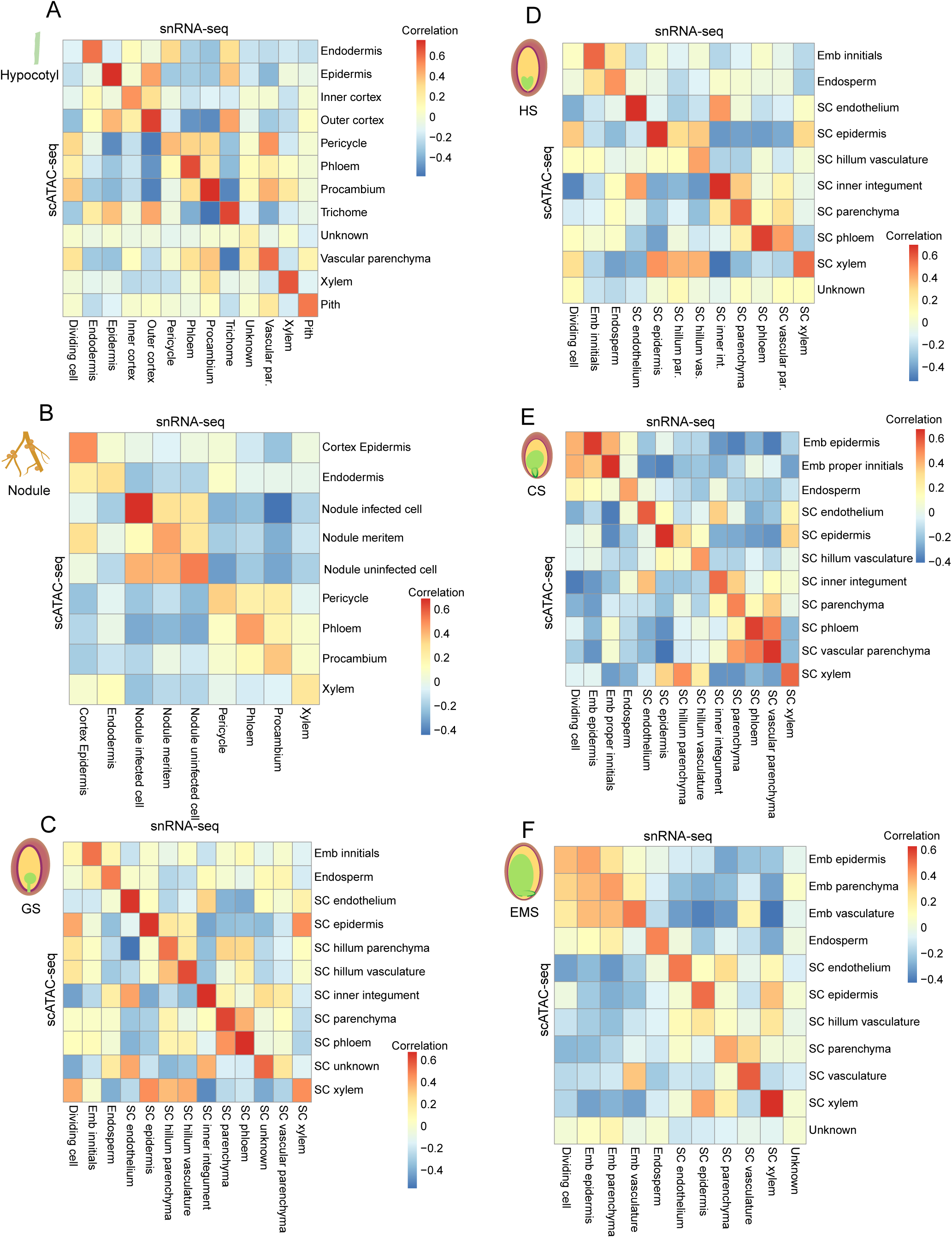
Gene expression or activity correlation between snRNA-seq and scATAC-seq, related to Figure 1. (A-F) The heatmap of spearman correlation coefficient between 1,000 most variable gene accessibility and expression across all cell types in each tissues, including hypocotyls (A), nodules (B), seeds at globular stage (C), heart stage (D), cotyledon stage (E) and early maturation stage (F).

**Figure S6.**
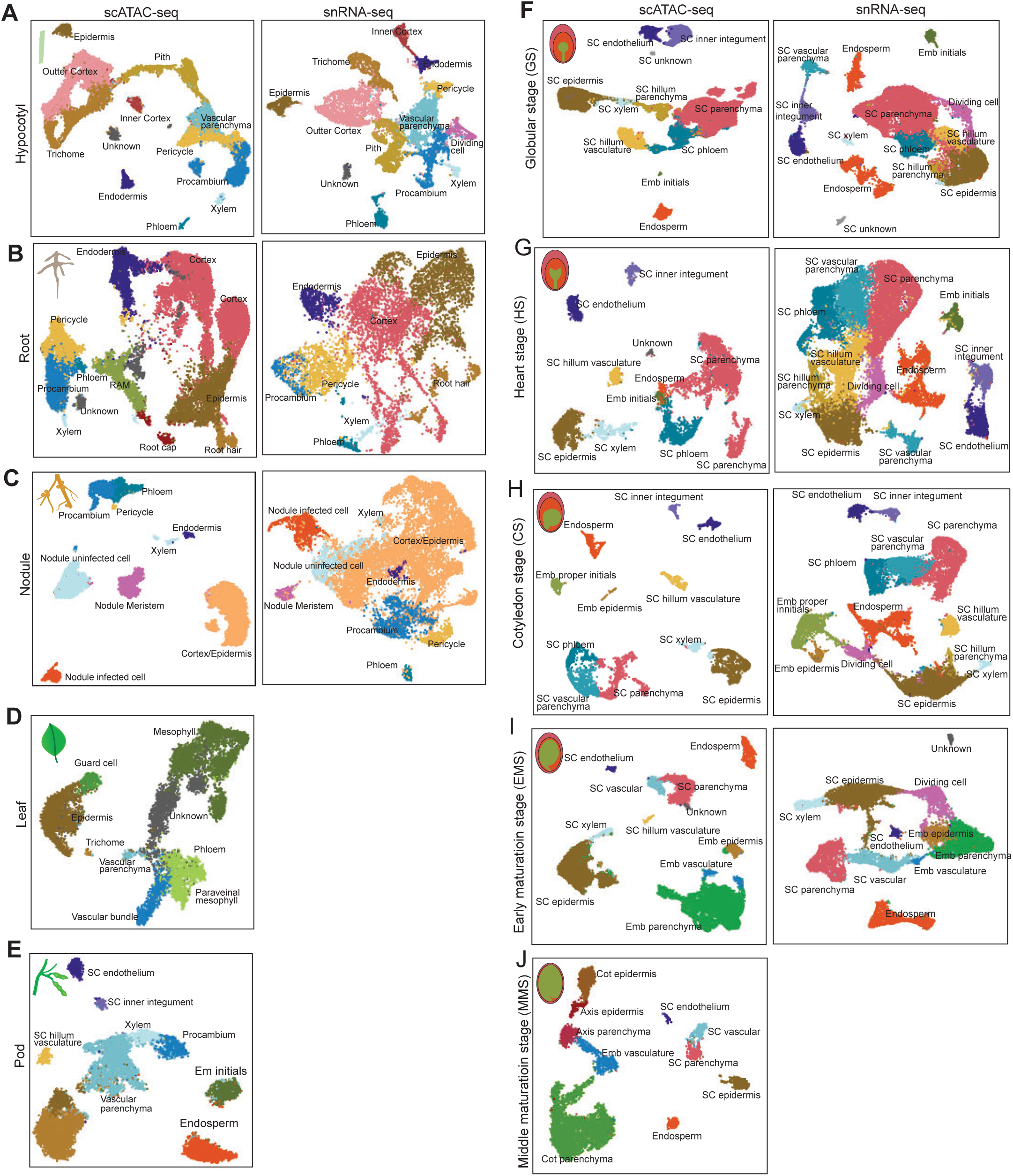
Cell type annotation for nuclei from scATAC-seq and snRNA-seq, related to Figure 1. (A-J) UMAP projection of nuclei, distinguished by assigned cell-type labels for scATAC-seq (left) snRNA-seq (right) across ten tissues, including hypocotyls (A), roots (B), nodules (C), leaves (D), pods (E), seeds at globular stage (F), heart stage (G), cotyledon stage (H), early maturation stage (I), and middle maturation stage (J).

**Figure S7.**
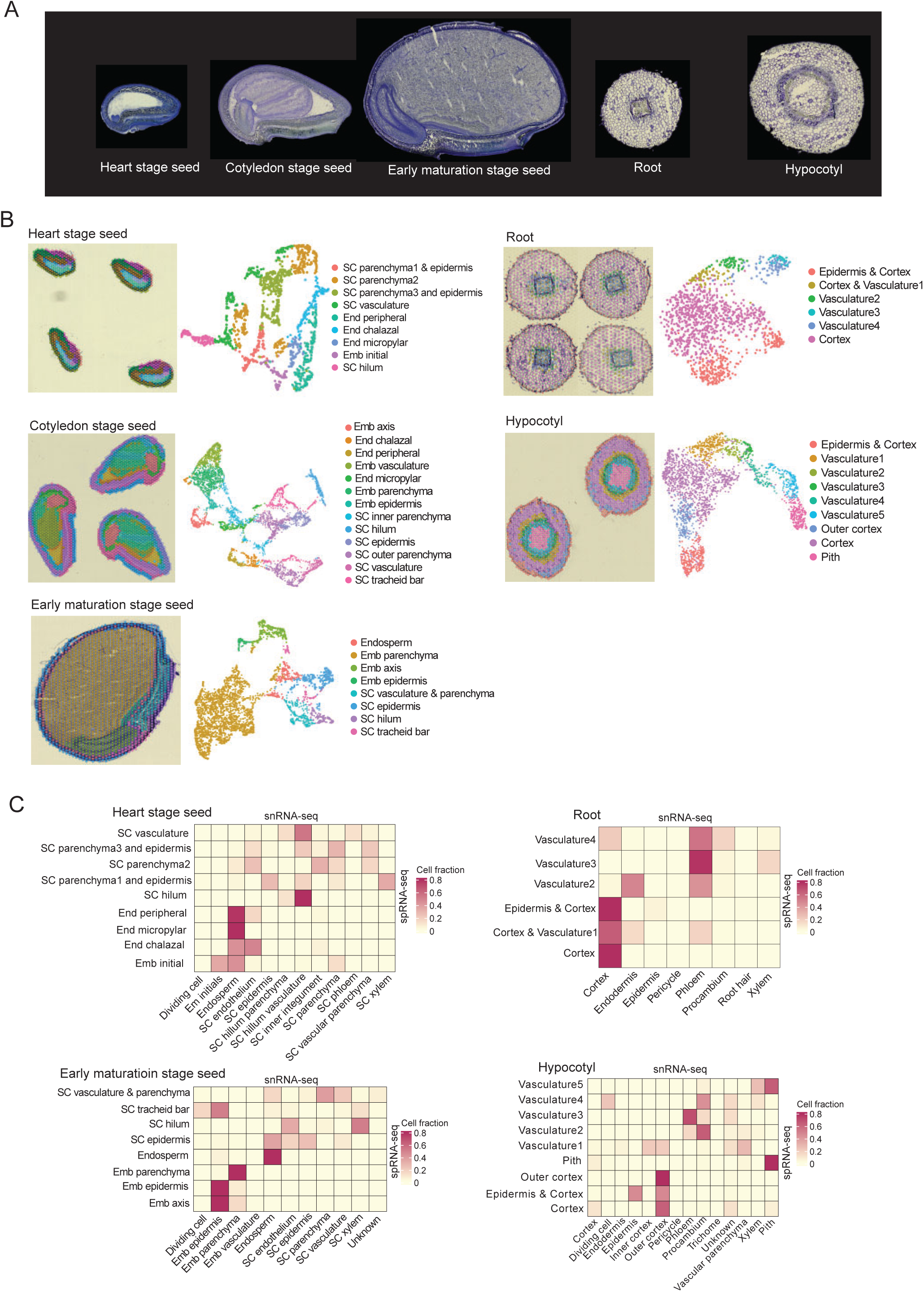
Spatial transcriptome atlas of soybean, related to Figure 2. (A) The histological structure of soybean tissues used for spRNA-seq. (B) The visualization of spatial spot clusters on the tissue (left) and on the UMAP plot (right) for all the tissue types. (C) Heatmaps of the snRNA-seq cell type prediction scores on the spRNA-seq cell types for all the tissue types.

**Figure S8.**
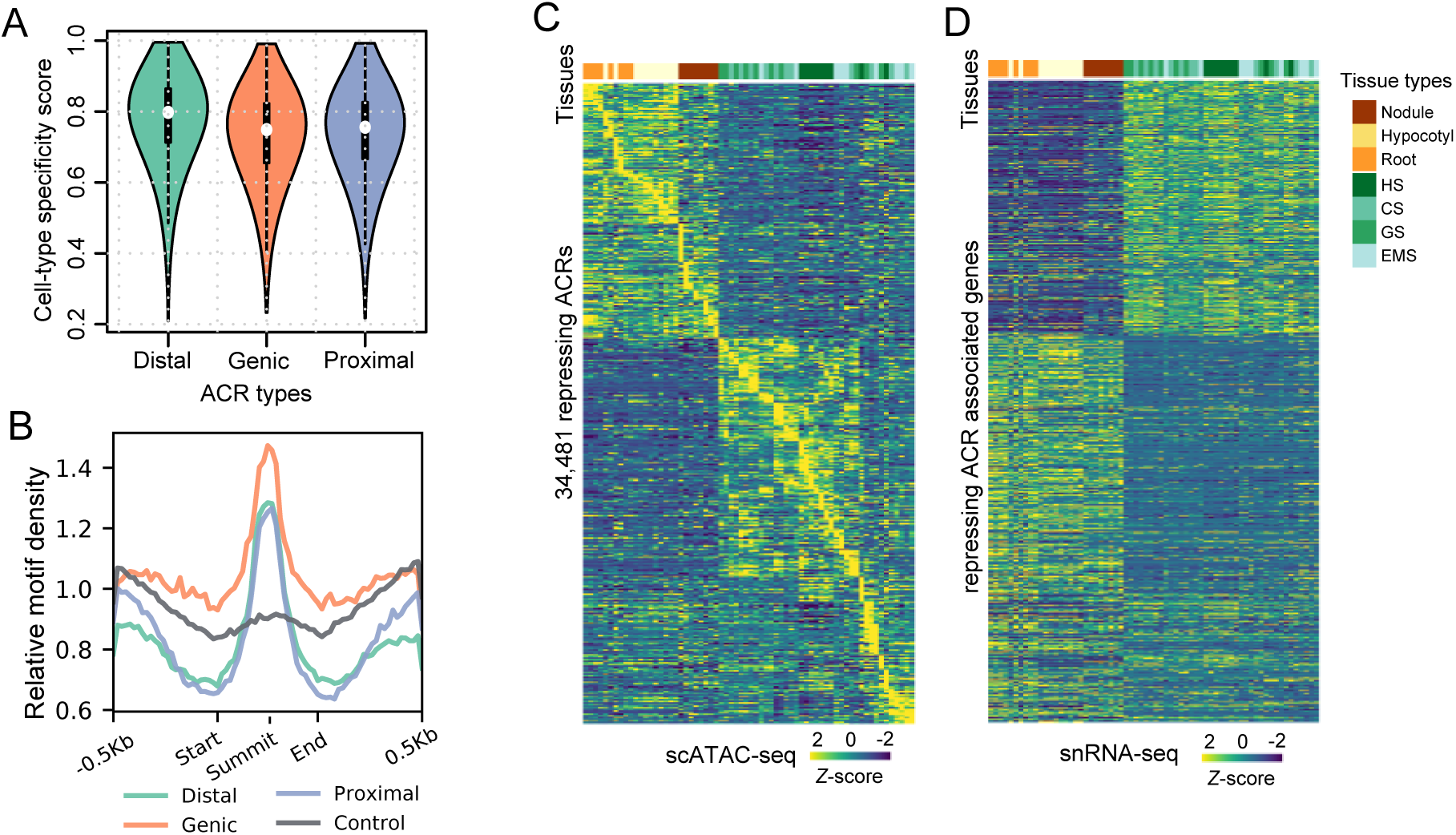
Characterization of ACRs, related to Figure 3. (A) Distribution of cell-type specificity score across three types of ACRs. (B) Relative density density within 500-bp flanking regions of different classes of ACRs and control regions. (C-D) Heatmap showing chromatin accessibility of repressing ACRs (C) and the expression of associated genes.

**Figure S9.**
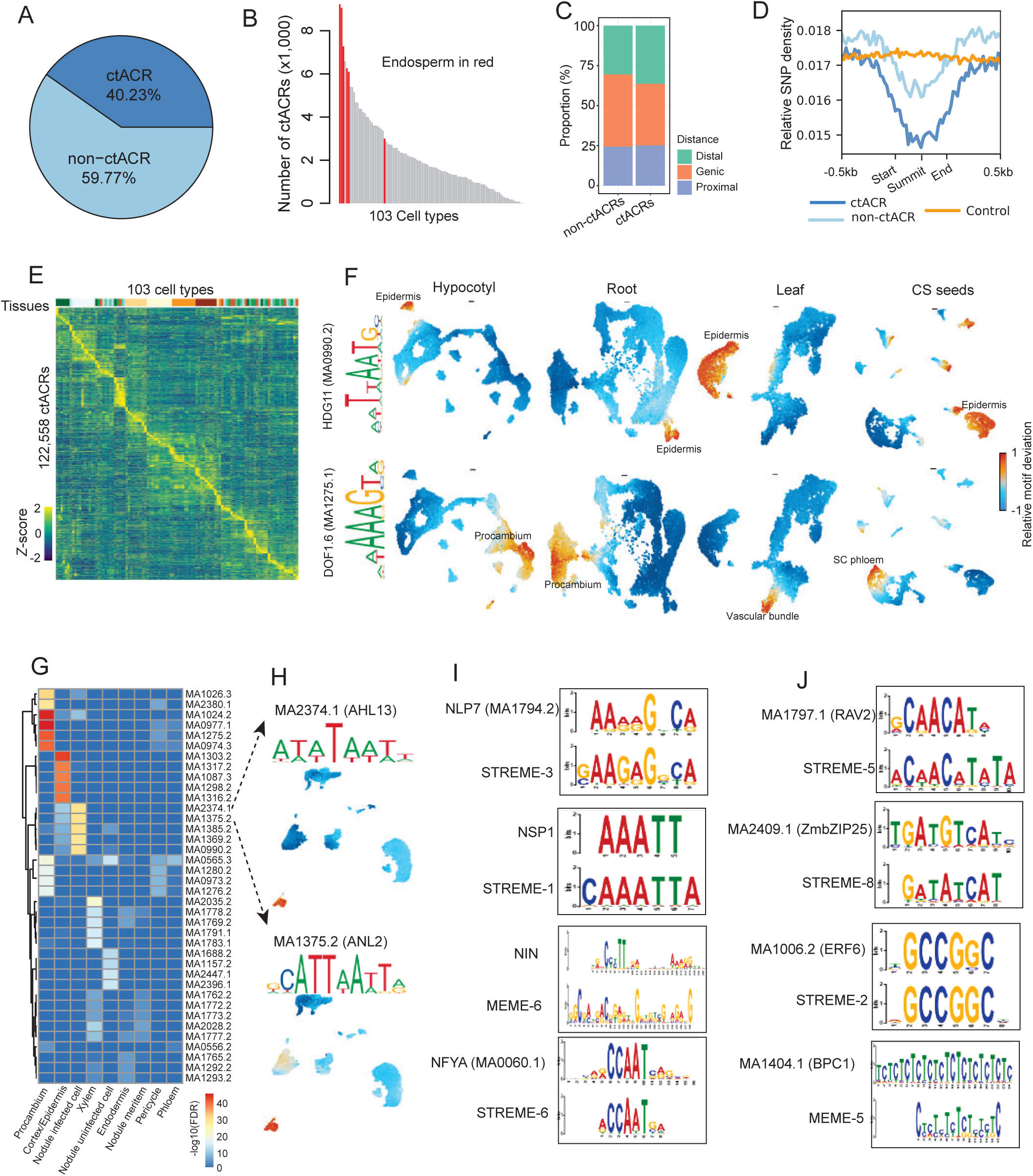
Characterization of ctACRs, related to Figure 4. (A) Proportion of ctACRs and non-ctACR. (B) Distribution of the number of ctACRs identified in each cell type. Endosperm cell types were highlighted in red. (C) Proportion of different groups of ACRs located in genic, proximal and distal regions. (D) Relative SNP density within 500-bp flanking regions of different groups of distal ACRs and control regions. (E) Heatmap showing relative chromatin accessibility of ctACRs across 103 cell types. (F) UMAP embeddings overlaid with motif deviation score of epidermis specific TF HDG11 (top row) and vasculature specific TF DOF1.6 (bottom row) across 4 tissues, including hypocotyls, roots, leaves, and seeds at cotyledon stage. (G) Heatmap of motif enrichment across 9 cell types in nodules. (H) UMAP embeddings overlaid with motif deviation score of motif MA2374.1 (top) and MA1375.2 (bottom) in nodule tissue. (I) The motif sequence alignment of key nodulation related TF motifs (up) and *de novo* motifs (bottom) enriched in infected-cell-specific ACRs. (J) The motif sequence alignment of known TF motifs in JASPAR2024 (up) and *de novo* motifs (bottom) enriched in infected-cell-specific ACRs.

**Figure S10.**
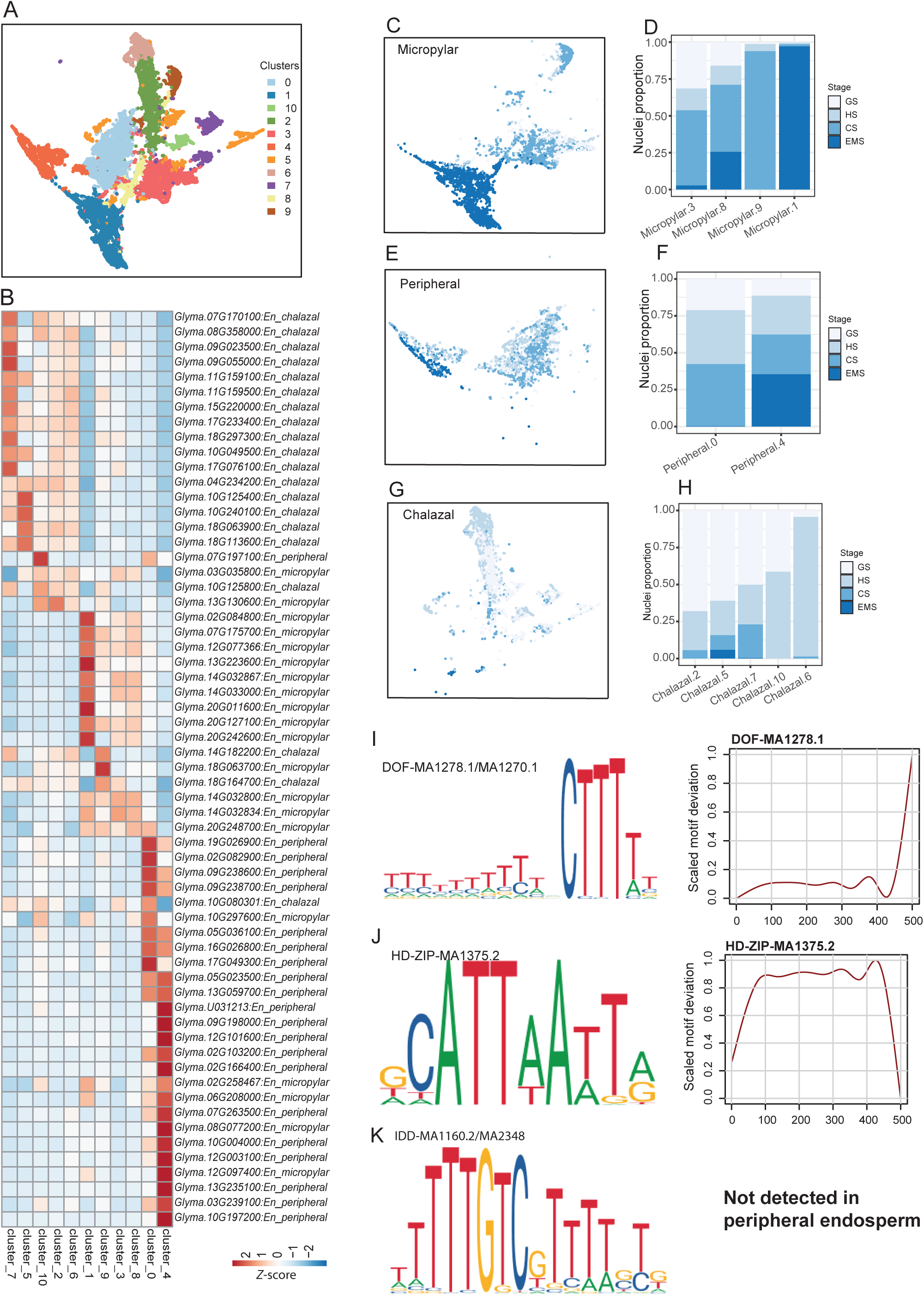
Characterizing three sub-cell types of endosperm, related to Figure 5. (A) UMAP embeddings of integration of scATAC-seq and snRNA-seq for endosperm cells across 4 developmental stages, including globular stage, heart stage, cotyledon stage and early maturation stage. (B) *Z*-score heatmap of gene expression for *de novo* marker genes for three sub-cell types of endosperm, including micropylar, peripheral and chalazal endosperm from spRNA-seq of seeds at the cotyledon stage. (C-D) UMAP embeddings of micropylar endosperm cells overlaid with four developmental stages (C) and nuclei proportion in four developmental stages across micropylar clusters (D). Seed stages include GS (globular stage), HS (heart stage), CS (cotyledon stage), EMS (early maturation stage). (E-F) Similar to panels C-D, but for the peripheral endosperm. (G-H) Similar to panels C-D, but for the chalazal endosperm. (I-K) The five motifs that were identified in ACRs of all the 13 SWEET transporter genes (left) and its motif deviation across peripheral endosperm developmental pseudotime (right).

**Figure S11.**
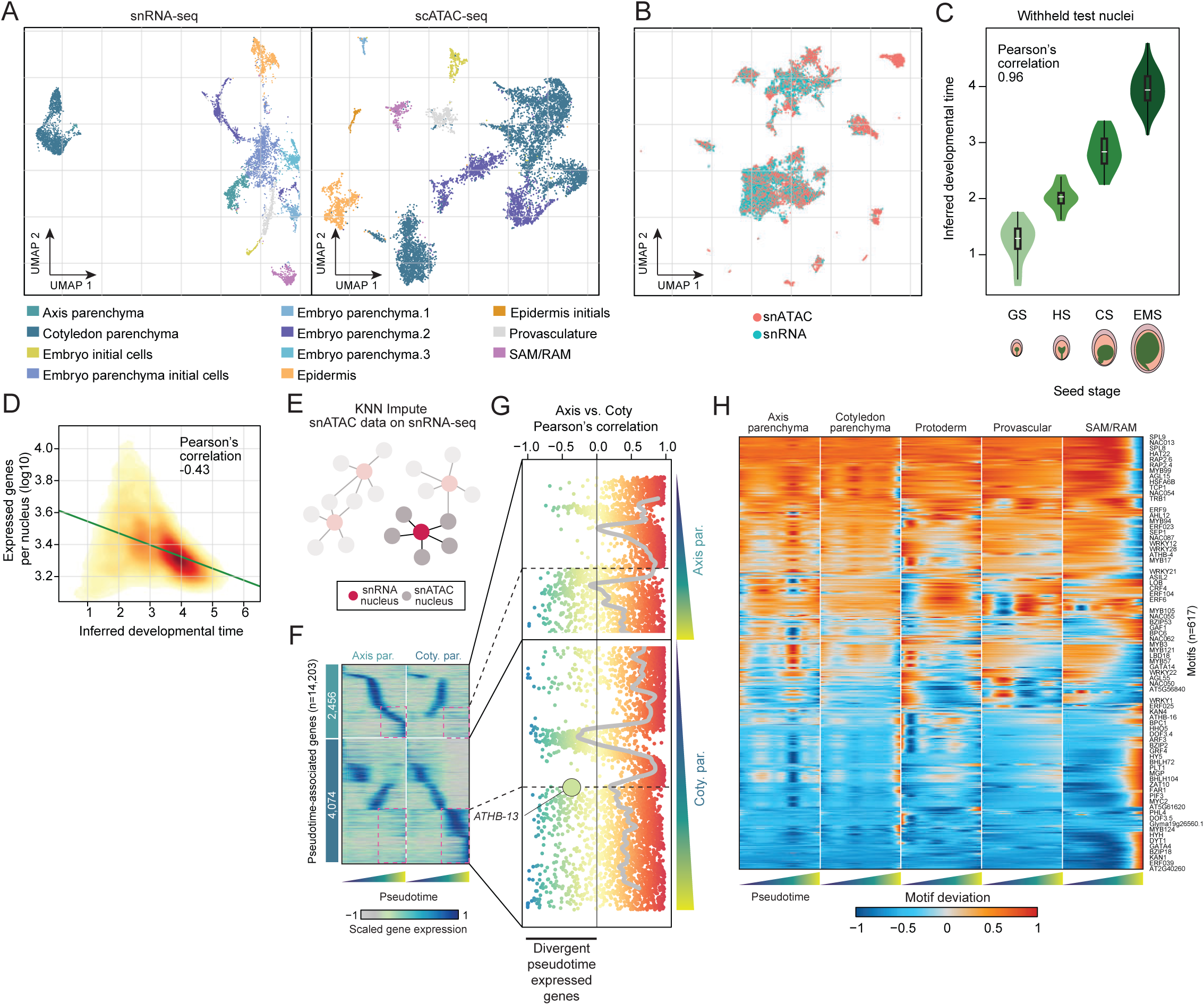
Analysis of embryogenesis trajectories, related to Figure 6. (A) Cell-type annotation of snRNA-seq and scATAC-seq embryogenic nuclei. (B) Integration of scATAC-seq and snRNA-seq embryo nuclei via non-negative matrix factorization. (C) Comparison of inferred nuclei age derived from LASSO predictions across seed developmental stages from withheld test nuclei. (D) Comparison of inferred nuclei age with the number of uniquely expressed genes (log10). (E) Illustration of scATAC-seq and snRNA-seq imputation strategy. (F) Gene expression dynamics across pseudotime for axis and cotyledon parenchyma trajectories. Red boxes highlight genes with divergent expression patterns. (G) Correlation of gene expression profiles between axis and cotyledon parenchyma trajectories. *ATHB-13* is highlighted. (H) TF motif deviation scores across pseudotime for the five embryogenesis branches.

**Figure S12.**
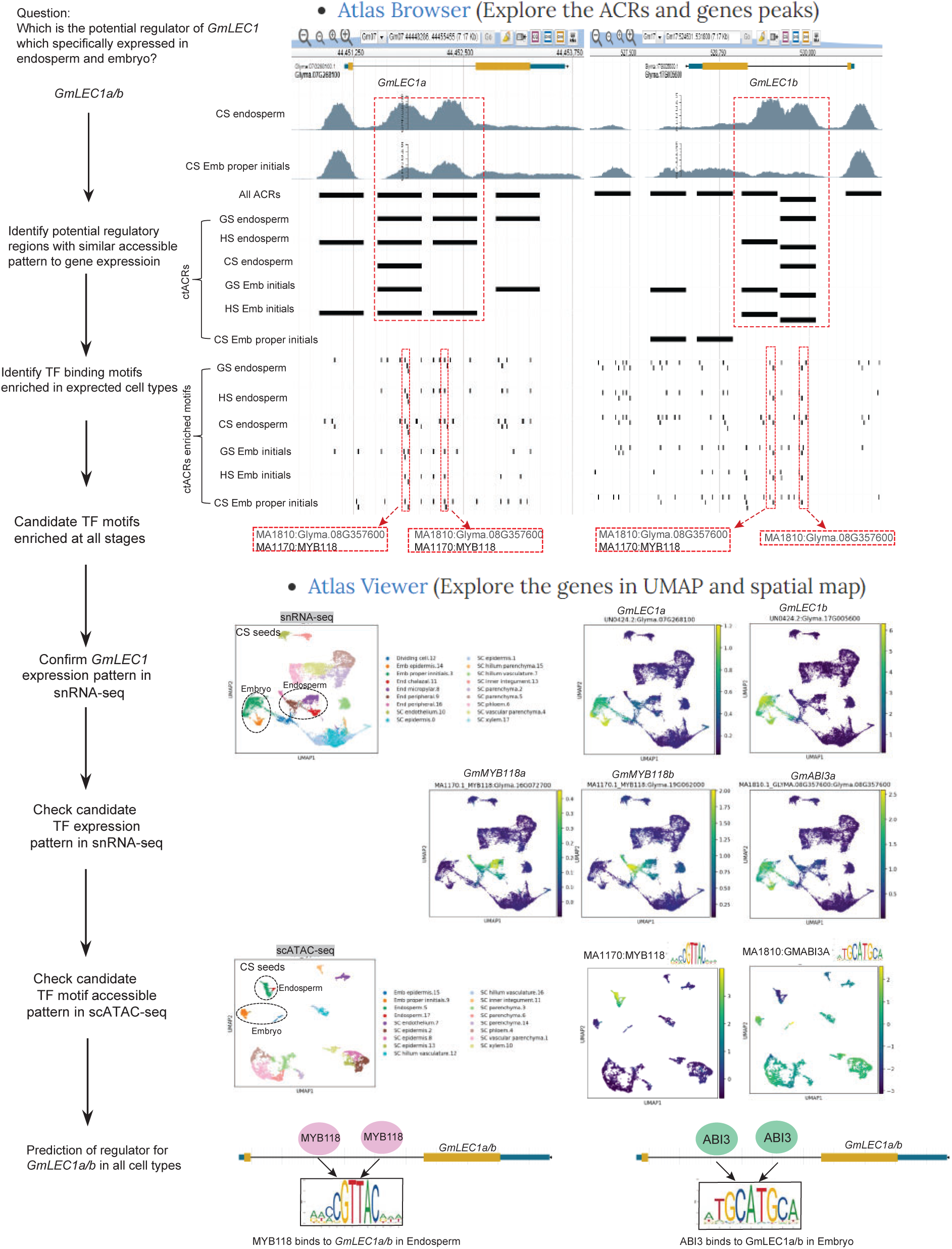
Workflow of exploring *GmLEC1a/b* gene regulatory network with soybean multi-omic atlas database.

## References

1. Schmitz, R.J., Grotewold, E., and Stam, M. (2022). Cis-regulatory sequences in plants: Their importance, discovery, and future challenges. Plant Cell 34, 718–741. 10.1093/plcell/koab281.

2. Marand, A.P., Eveland, A.L., Kaufmann, K., and Springer, N.M. (2023). cis-Regulatory Elements in Plant Development, Adaptation, and Evolution. Annu Rev Plant Biol 74, 111–137. 10.1146/annurev-arplant-070122-030236.

3. Klemm, S.L., Shipony, Z., and Greenleaf, W.J. (2019). Chromatin accessibility and the regulatory epigenome. Nat Rev Genet 20, 207–220. 10.1038/s41576-018-0089-8.

4. Vandereyken, K., Sifrim, A., Thienpont, B., and Voet, T. (2023). Methods and applications for single-cell and spatial multi-omics. Nat Rev Genet 24, 494–515. 10.1038/s41576-023-00580-2.

5. Zhang, X., Marand, A.P., Yan, H., and Schmitz, R.J. (2024). scifi-ATAC-seq: massive-scale single-cell chromatin accessibility sequencing using combinatorial fluidic indexing. Genome Biol 25, 90. 10.1186/s13059-024-03235-5.

6. Hie, B., Peters, J., Nyquist, S.K., Shalek, A.K., Berger, B., and Bryson, B.D. (2020). Computational methods for single-cell RNA sequencing. Annual Review of Biomedical Data Science 3, 339–364.

7. Cao, J., O’Day, D.R., Pliner, H.A., Kingsley, P.D., Deng, M., Daza, R.M., Zager, M.A., Aldinger, K.A., Blecher-Gonen, R., Zhang, F., et al. (2020). A human cell atlas of fetal gene expression. Science 370. 10.1126/science.aba7721.

8. Zhang, K., Hocker, J.D., Miller, M., Hou, X., Chiou, J., Poirion, O.B., Qiu, Y., Li, Y.E., Gaulton, K.J., Wang, A., et al. (2021). A single-cell atlas of chromatin accessibility in the human genome. Cell 184, 5985–6001 e5919. 10.1016/j.cell.2021.10.024.

9. Yao, Z., Liu, H., Xie, F., Fischer, S., Adkins, R.S., Aldridge, A.I., Ament, S.A., Bartlett, A., Behrens, M.M., Van den Berge, K., et al. (2021). A transcriptomic and epigenomic cell atlas of the mouse primary motor cortex. Nature 598, 103–110. 10.1038/s41586-021-03500-8.

10. Li, Y.E., Preissl, S., Miller, M., Johnson, N.D., Wang, Z., Jiao, H., Zhu, C., Wang, Z., Xie, Y., Poirion, O., et al. (2023). A comparative atlas of single-cell chromatin accessibility in the human brain. Science 382, eadf7044. 10.1126/science.adf7044.

11. Cuperus, J.T. (2022). Single-cell genomics in plants: current state, future directions, and hurdles to overcome. Plant Physiol 188, 749–755. 10.1093/plphys/kiab478.

12. Shaw, R., Tian, X., and Xu, J. (2021). Single-Cell Transcriptome Analysis in Plants: Advances and Challenges. Mol Plant 14, 115–126. 10.1016/j.molp.2020.10.012.

13. Bang, S., Zhang, X., Gregory, J., Chen, Z., Galli, M., Gallavotti, A., and Schmitz, R.J. (2024). WUSCHEL dependent chromatin regulation in maize inflorescence development at single-cell resolution. bioRxiv, 2024.2005.2013.593957. 10.1101/2024.05.13.593957.

14. Mendieta, J.P., Tu, X., Jiang, D., Yan, H., Zhang, X., Marand, A.P., Zhong, S., and Schmitz, R.J. (2024). Investigating the cis-Regulatory Basis of C3 and C4 Photosynthesis in Grasses at Single-Cell Resolution. bioRxiv, 2024.2001.2005.574340. 10.1101/2024.01.05.574340.

15. Farmer, A., Thibivilliers, S., Ryu, K.H., Schiefelbein, J., and Libault, M. (2021). Single-nucleus RNA and ATAC sequencing reveals the impact of chromatin accessibility on gene expression in Arabidopsis roots at the single-cell level. Mol Plant 14, 372–383. 10.1016/j.molp.2021.01.001.

16. Shahan, R., Hsu, C.W., Nolan, T.M., Cole, B.J., Taylor, I.W., Greenstreet, L., Zhang, S., Afanassiev, A., Vlot, A.H.C., Schiebinger, G., et al. (2022). A single-cell Arabidopsis root atlas reveals developmental trajectories in wild-type and cell identity mutants. Dev Cell 57, 543–560 e549. 10.1016/j.devcel.2022.01.008.

17. Xu, X., Crow, M., Rice, B.R., Li, F., Harris, B., Liu, L., Demesa-Arevalo, E., Lu, Z., Wang, L., Fox, N., et al. (2021). Single-cell RNA sequencing of developing maize ears facilitates functional analysis and trait candidate gene discovery. Dev Cell 56, 557–568 e556. 10.1016/j.devcel.2020.12.015.

18. Lee, T.A., Nobori, T., Illouz-Eliaz, N., Xu, J., Jow, B., Nery, J.R., and Ecker, J.R. (2023). A single-nucleus atlas of seed-to-seed development in Arabidopsis. bioRxiv, 2023.2003. 2023.533992.

19. Marand, A.P., Chen, Z., Gallavotti, A., and Schmitz, R.J. (2021). A cis-regulatory atlas in maize at single-cell resolution. Cell 184, 3041–3055 e3021. 10.1016/j.cell.2021.04.014.

20. Yan, H., Mendieta, J.P., Zhang, X., Marand, A.P., Liang, Y., Luo, Z., Minow, M.A.A., Roule, T., Wagner, D., Tu, X., et al. (2024). Evolution of plant cell-type-specific cis -regulatory elements. bioRxiv. 10.1101/2024.01.08.574753.

21. Mendieta, J.P., Sangra, A., Yan, H., Minow, M.A.A., and Schmitz, R.J. (2023). Exploring plant cis-regulatory elements at single-cell resolution: overcoming biological and computational challenges to advance plant research. Plant J 115, 1486–1499. 10.1111/tpj.16351.

22. Liu, Z., Kong, X., Long, Y., Liu, S., Zhang, H., Jia, J., Cui, W., Zhang, Z., Song, X., Qiu, L., et al. (2023). Integrated single-nucleus and spatial transcriptomics captures transitional states in soybean nodule maturation. Nat Plants 9, 515–524. 10.1038/s41477-023-01387-z.

23. Yu, X., Liu, Z., and Sun, X. (2023). Single-cell and spatial multi-omics in the plant sciences: Technical advances, applications, and perspectives. Plant Commun 4, 100508. 10.1016/j.xplc.2022.100508.

24. Wang, Y., Luo, Y., Guo, X., Li, Y., Yan, J., Shao, W., Wei, W., Wei, X., Yang, T., Chen, J., et al. (2024). A spatial transcriptome map of the developing maize ear. Nat Plants 10, 815–827. 10.1038/s41477-024-01683-2.

25. Schmutz, J., Cannon, S.B., Schlueter, J., Ma, J., Mitros, T., Nelson, W., Hyten, D.L., Song, Q., Thelen, J.J., Cheng, J., et al. (2010). Genome sequence of the palaeopolyploid soybean. Nature 463, 178–183. 10.1038/nature08670.

26. Hao, Y., Hao, S., Andersen-Nissen, E., Mauck, W.M., 3rd, Zheng, S., Butler, A., Lee, M.J., Wilk, A.J., Darby, C., Zager, M., et al. (2021). Integrated analysis of multimodal single-cell data. Cell 184, 3573–3587 e3529. 10.1016/j.cell.2021.04.048.

27. Danzer, J., Mellott, E., Bui, A.Q., Le, B.H., Martin, P., Hashimoto, M., Perez-Lesher, J., Chen, M., Pelletier, J.M., Somers, D.A., et al. (2015). Down-Regulating the Expression of 53 Soybean Transcription Factor Genes Uncovers a Role for SPEECHLESS in Initiating Stomatal Cell Lineages during Embryo Development. Plant Physiol 168, 1025–1035. 10.1104/pp.15.00432.

28. Jo, L., Pelletier, J.M., Hsu, S.W., Baden, R., Goldberg, R.B., and Harada, J.J. (2020). Combinatorial interactions of the LEC1 transcription factor specify diverse developmental programs during soybean seed development. Proc Natl Acad Sci U S A 117, 1223–1232. 10.1073/pnas.1918441117.

29. Wang, S., Yokosho, K., Guo, R., Whelan, J., Ruan, Y.L., Ma, J.F., and Shou, H. (2019). The Soybean Sugar Transporter GmSWEET15 Mediates Sucrose Export from Endosperm to Early Embryo. Plant Physiol 180, 2133–2141. 10.1104/pp.19.00641.

30. Perez-Grau, L., and Goldberg, R.B. (1989). Soybean Seed Protein Genes Are Regulated Spatially during Embryogenesis. Plant Cell 1, 1095–1109. 10.1105/tpc.1.11.1095.

31. Aida, M., Beis, D., Heidstra, R., Willemsen, V., Blilou, I., Galinha, C., Nussaume, L., Noh, Y.S., Amasino, R., and Scheres, B. (2004). The PLETHORA genes mediate patterning of the Arabidopsis root stem cell niche. Cell 119, 109–120. 10.1016/j.cell.2004.09.018.

32. Wang, S., Liu, S., Wang, J., Yokosho, K., Zhou, B., Yu, Y.C., Liu, Z., Frommer, W.B., Ma, J.F., Chen, L.Q., et al. (2020). Simultaneous changes in seed size, oil content and protein content driven by selection of SWEET homologues during soybean domestication. Natl Sci Rev 7, 1776–1786. 10.1093/nsr/nwaa110.

33. Torkamaneh, D., Laroche, J., Valliyodan, B., O’Donoughue, L., Cober, E., Rajcan, I., Vilela Abdelnoor, R., Sreedasyam, A., Schmutz, J., Nguyen, H.T., and Belzile, F. (2021). Soybean (Glycine max) Haplotype Map (GmHapMap): a universal resource for soybean translational and functional genomics. Plant Biotechnol J 19, 324–334. 10.1111/pbi.13466.

34. Marand, A.P., and Schmitz, R.J. (2022). Single-cell analysis of cis-regulatory elements. Curr Opin Plant Biol 65, 102094. 10.1016/j.pbi.2021.102094.

35. Fang, C., Yang, M., Tang, Y., Zhang, L., Zhao, H., Ni, H., Chen, Q., Meng, F., and Jiang, J. (2023). Dynamics of cis-regulatory sequences and transcriptional divergence of duplicated genes in soybean. Proc Natl Acad Sci U S A 120, e2303836120. 10.1073/pnas.2303836120.

36. Soyano, T., Shimoda, Y., Kawaguchi, M., and Hayashi, M. (2019). A shared gene drives lateral root development and root nodule symbiosis pathways in Lotus. Science 366, 1021–1023. 10.1126/science.aax2153.

37. Dong, Y., Yang, X., Liu, J., Wang, B.H., Liu, B.L., and Wang, Y.Z. (2014). Pod shattering resistance associated with domestication is mediated by a NAC gene in soybean. Nat Commun 5, 3352. 10.1038/ncomms4352.

38. Rauf, M., Arif, M., Fisahn, J., Xue, G.P., Balazadeh, S., and Mueller-Roeber, B. (2013). NAC transcription factor speedy hyponastic growth regulates flooding-induced leaf movement in Arabidopsis. Plant Cell 25, 4941–4955. 10.1105/tpc.113.117861.

39. Liu, Y., Peng, X., Ma, A., Liu, W., Liu, B., Yun, D.J., and Xu, Z.Y. (2023). Type-B response regulator OsRR22 forms a transcriptional activation complex with OsSLR1 to modulate OsHKT2;1 expression in rice. Sci China Life Sci 66, 2922–2934. 10.1007/s11427-023-2464-2.

40. Mara, C.D., Huang, T., and Irish, V.F. (2010). The Arabidopsis floral homeotic proteins APETALA3 and PISTILLATA negatively regulate the BANQUO genes implicated in light signaling. Plant Cell 22, 690–702. 10.1105/tpc.109.065946.

41. Zhao, P.X., Zhang, J., Chen, S.Y., Wu, J., Xia, J.Q., Sun, L.Q., Ma, S.S., and Xiang, C.B. (2021). Arabidopsis MADS-box factor AGL16 is a negative regulator of plant response to salt stress by downregulating salt-responsive genes. New Phytol 232, 2418–2439. 10.1111/nph.17760.

42. Ng, M., and Yanofsky, M.F. (2001). Function and evolution of the plant MADS-box gene family. Nature Reviews Genetics 2, 186–195.

43. Fueyo, R., Judd, J., Feschotte, C., and Wysocka, J. (2022). Roles of transposable elements in the regulation of mammalian transcription. Nature reviews Molecular cell biology 23, 481–497.

44. Ricci, W.A., Lu, Z., Ji, L., Marand, A.P., Ethridge, C.L., Murphy, N.G., Noshay, J.M., Galli, M., Mejia-Guerra, M.K., Colome-Tatche, M., et al. (2019). Widespread long-range cis-regulatory elements in the maize genome. Nat Plants 5, 1237–1249. 10.1038/s41477-019-0547-0.

45. Rauluseviciute, I., Riudavets-Puig, R., Blanc-Mathieu, R., Castro-Mondragon, J.A., Ferenc, K., Kumar, V., Lemma, R.B., Lucas, J., Cheneby, J., Baranasic, D., et al. (2024). JASPAR 2024: 20th anniversary of the open-access database of transcription factor binding profiles. Nucleic Acids Res 52, D174–D182. 10.1093/nar/gkad1059.

46. Khosla, A., Paper, J.M., Boehler, A.P., Bradley, A.M., Neumann, T.R., and Schrick, K. (2014). HD-Zip Proteins GL2 and HDG11 Have Redundant Functions in Arabidopsis Trichomes, and GL2 Activates a Positive Feedback Loop via MYB23. Plant Cell 26, 2184–2200. 10.1105/tpc.113.120360.

47. Barthole, G., To, A., Marchive, C., Brunaud, V., Soubigou-Taconnat, L., Berger, N., Dubreucq, B., Lepiniec, L., and Baud, S. (2014). MYB118 represses endosperm maturation in seeds of Arabidopsis. Plant Cell 26, 3519–3537. 10.1105/tpc.114.130021.

48. Roy, S., Liu, W., Nandety, R.S., Crook, A., Mysore, K.S., Pislariu, C.I., Frugoli, J., Dickstein, R., and Udvardi, M.K. (2020). Celebrating 20 Years of Genetic Discoveries in Legume Nodulation and Symbiotic Nitrogen Fixation. Plant Cell 32, 15–41. 10.1105/tpc.19.00279.

49. Wu, X., Xiong, Y., Lu, J., Yang, M., Ji, H., Li, X., and Wang, Z. (2023). GmNLP7a inhibits soybean nodulation by interacting with GmNIN1a. The Crop Journal 11, 1401–1410.

50. Nishida, H., Nosaki, S., Suzuki, T., Ito, M., Miyakawa, T., Nomoto, M., Tada, Y., Miura, K., Tanokura, M., Kawaguchi, M., and Suzaki, T. (2021). Different DNA-binding specificities of NLP and NIN transcription factors underlie nitrate-induced control of root nodulation. Plant Cell 33, 2340–2359. 10.1093/plcell/koab103.

51. Zhao, J., Favero, D.S., Peng, H., and Neff, M.M. (2013). Arabidopsis thaliana AHL family modulates hypocotyl growth redundantly by interacting with each other via the PPC/DUF296 domain. Proc Natl Acad Sci U S A 110, E4688–4697. 10.1073/pnas.1219277110.

52. Kubo, H., Peeters, A.J., Aarts, M.G., Pereira, A., and Koornneef, M. (1999). ANTHOCYANINLESS2, a homeobox gene affecting anthocyanin distribution and root development in Arabidopsis. Plant Cell 11, 1217–1226. 10.1105/tpc.11.7.1217.

53. Andriankaja, A., Boisson-Dernier, A., Frances, L., Sauviac, L., Jauneau, A., Barker, D.G., and de Carvalho-Niebel, F. (2007). AP2-ERF transcription factors mediate Nod factor dependent Mt ENOD11 activation in root hairs via a novel cis-regulatory motif. Plant Cell 19, 2866–2885. 10.1105/tpc.107.052944.

54. Doll, N.M., and Ingram, G.C. (2022). Embryo–endosperm interactions. Annual Review of Plant Biology 73, 293–321.

55. Povilus, R.A., and Gehring, M. (2022). Maternal-filial transfer structures in endosperm: A nexus of nutritional dynamics and seed development. Curr Opin Plant Biol 65, 102121. 10.1016/j.pbi.2021.102121.

56. Picard, C.L., Povilus, R.A., Williams, B.P., and Gehring, M. (2021). Transcriptional and imprinting complexity in Arabidopsis seeds at single-nucleus resolution. Nat Plants 7, 730–738. 10.1038/s41477-021-00922-0.

57. Dute, R.R., and Peterson, C.M. (1992). Early Endosperm Development in Ovules of Soybean, Glycine max (L) Merr. (Fabaceae)*. Annals of Botany 69, 263–271. 10.1093/oxfordjournals.aob.a088339.

58. Belmonte, M.F., Kirkbride, R.C., Stone, S.L., Pelletier, J.M., Bui, A.Q., Yeung, E.C., Hashimoto, M., Fei, J., Harada, C.M., Munoz, M.D., et al. (2013). Comprehensive developmental profiles of gene activity in regions and subregions of the Arabidopsis seed. Proc Natl Acad Sci U S A 110, E435–444. 10.1073/pnas.1222061110.

59. Nguyen, H., Brown, R., and Lemmon, B. (2000). The specialized chalazal endosperm in Arabidopsis thaliana and Lepidium virginicum (Brassicaceae). Protoplasma 212, 99–110.

60. Doll, N.M., Royek, S., Fujita, S., Okuda, S., Chamot, S., Stintzi, A., Widiez, T., Hothorn, M., Schaller, A., Geldner, N., and Ingram, G. (2020). A two-way molecular dialogue between embryo and endosperm is required for seed development. Science 367, 431–435. 10.1126/science.aaz4131.

61. Doll, N.M., and Nowack, M.K. (2024). Endosperm cell death: roles and regulation in angiosperms. J Exp Bot. 10.1093/jxb/erae052.

62. Xiong, H., Wang, W., and Sun, M.X. (2021). Endosperm development is an autonomously programmed process independent of embryogenesis. Plant Cell 33, 1151–1160. 10.1093/plcell/koab007.

63. Buono, R.A., Hudecek, R., and Nowack, M.K. (2019). Plant proteases during developmental programmed cell death. J Exp Bot 70, 2097–2112. 10.1093/jxb/erz072.

64. Patil, G., Valliyodan, B., Deshmukh, R., Prince, S., Nicander, B., Zhao, M., Sonah, H., Song, L., Lin, L., Chaudhary, J., et al. (2015). Soybean (Glycine max) SWEET gene family: insights through comparative genomics, transcriptome profiling and whole genome re-sequence analysis. BMC Genomics 16, 520. 10.1186/s12864-015-1730-y.

65. Braun, D.M. (2022). Phloem Loading and Unloading of Sucrose: What a Long, Strange Trip from Source to Sink. Annu Rev Plant Biol 73, 553–584. 10.1146/annurev-arplant-070721-083240.

66. Julius, B.T., Leach, K.A., Tran, T.M., Mertz, R.A., and Braun, D.M. (2017). Sugar Transporters in Plants: New Insights and Discoveries. Plant Cell Physiol 58, 1442–1460. 10.1093/pcp/pcx090.

67. Wang, H.W., Zhang, B., Hao, Y.J., Huang, J., Tian, A.G., Liao, Y., Zhang, J.S., and Chen, S.Y. (2007). The soybean Dof-type transcription factor genes, GmDof4 and GmDof11, enhance lipid content in the seeds of transgenic Arabidopsis plants. Plant J 52, 716–729. 10.1111/j.1365-313X.2007.03268.x.

68. Calderon, D., Blecher-Gonen, R., Huang, X., Secchia, S., Kentro, J., Daza, R.M., Martin, B., Dulja, A., Schaub, C., Trapnell, C., et al. (2022). The continuum of Drosophila embryonic development at single-cell resolution. Science 377, eabn5800. 10.1126/science.abn5800.

69. Candat, A., Paszkiewicz, G., Neveu, M., Gautier, R., Logan, D.C., Avelange-Macherel, M.H., and Macherel, D. (2014). The ubiquitous distribution of late embryogenesis abundant proteins across cell compartments in Arabidopsis offers tailored protection against abiotic stress. Plant Cell 26, 3148–3166. 10.1105/tpc.114.127316.

70. ten Hove, C.A., Lu, K.J., and Weijers, D. (2015). Building a plant: cell fate specification in the early Arabidopsis embryo. Development 142, 420–430. 10.1242/dev.111500.

71. Thien Nguyen, Q., Kisiala, A., Andreas, P., Neil Emery, R., and Narine, S. (2016). Soybean seed development: fatty acid and phytohormone metabolism and their interactions. Current Genomics 17, 241–260.

72. Le, B.H., Cheng, C., Bui, A.Q., Wagmaister, J.A., Henry, K.F., Pelletier, J., Kwong, L., Belmonte, M., Kirkbride, R., Horvath, S., et al. (2010). Global analysis of gene activity during Arabidopsis seed development and identification of seed-specific transcription factors. Proc Natl Acad Sci U S A 107, 8063–8070. 10.1073/pnas.1003530107.

73. Silva, A.T., Ribone, P.A., Chan, R.L., Ligterink, W., and Hilhorst, H.W. (2016). A Predictive Coexpression Network Identifies Novel Genes Controlling the Seed-to-Seedling Phase Transition in Arabidopsis thaliana. Plant Physiol 170, 2218–2231. 10.1104/pp.15.01704.

74. Purugganan, M.D., and Jackson, S.A. (2021). Advancing crop genomics from lab to field. Nat Genet 53, 595–601. 10.1038/s41588-021-00866-3.

75. Wu, Y., Lee, S.K., Yoo, Y., Wei, J., Kwon, S.Y., Lee, S.W., Jeon, J.S., and An, G. (2018). Rice Transcription Factor OsDOF11 Modulates Sugar Transport by Promoting Expression of Sucrose Transporter and SWEET Genes. Mol Plant 11, 833–845. 10.1016/j.molp.2018.04.002.

76. Egli, D.B. (2017). Seed biology and yield of grain crops (CABI).

77. Chen, M., Lin, J.Y., Hur, J., Pelletier, J.M., Baden, R., Pellegrini, M., Harada, J.J., and Goldberg, R.B. (2018). Seed genome hypomethylated regions are enriched in transcription factor genes. Proc Natl Acad Sci U S A 115, E8315–E8322. 10.1073/pnas.1811017115.

78. Jo, L., Pelletier, J.M., and Harada, J.J. (2019). Central role of the LEAFY COTYLEDON1 transcription factor in seed development. J Integr Plant Biol 61, 564–580. 10.1111/jipb.12806.

79. Sayers, E.W., Beck, J., Bolton, E.E., Brister, J.R., Chan, J., Comeau, D.C., Connor, R., DiCuccio, M., Farrell, C.M., Feldgarden, M., et al. (2024). Database resources of the National Center for Biotechnology Information. Nucleic Acids Res 52, D33–D43. 10.1093/nar/gkad1044.

80. Brown, A.V., Conners, S.I., Huang, W., Wilkey, A.P., Grant, D., Weeks, N.T., Cannon, S.B., Graham, M.A., and Nelson, R.T. (2021). A new decade and new data at SoyBase, the USDA-ARS soybean genetics and genomics database. Nucleic Acids Res 49, D1496–D1501. 10.1093/nar/gkaa1107.

81. Roy Choudhury, S., Johns, S.M., and Pandey, S. (2019). A convenient, soil-free method for the production of root nodules in soybean to study the effects of exogenous additives. Plant Direct 3, e00135. 10.1002/pld3.135.

82. Pelletier, J.M., Kwong, R.W., Park, S., Le, B.H., Baden, R., Cagliari, A., Hashimoto, M., Munoz, M.D., Fischer, R.L., Goldberg, R.B., and Harada, J.J. (2017). LEC1 sequentially regulates the transcription of genes involved in diverse developmental processes during seed development. Proc Natl Acad Sci U S A 114, E6710–E6719. 10.1073/pnas.1707957114.

83. Lu, Z., Marand, A.P., Ricci, W.A., Ethridge, C.L., Zhang, X., and Schmitz, R.J. (2019). The prevalence, evolution and chromatin signatures of plant regulatory elements. Nat Plants 5, 1250–1259. 10.1038/s41477-019-0548-z.

84. Valliyodan, B., Cannon, S.B., Bayer, P.E., Shu, S., Brown, A.V., Ren, L., Jenkins, J., Chung, C.Y., Chan, T.F., Daum, C.G., et al. (2019). Construction and comparison of three reference-quality genome assemblies for soybean. Plant J 100, 1066–1082. 10.1111/tpj.14500.

85. Danecek, P., Bonfield, J.K., Liddle, J., Marshall, J., Ohan, V., Pollard, M.O., Whitwham, A., Keane, T., McCarthy, S.A., Davies, R.M., and Li, H. (2021). Twelve years of SAMtools and BCFtools. Gigascience 10. 10.1093/gigascience/giab008.

86. Blibaum, A., Werner, J., and Dobin, A. (2019). STARsolo: single-cell RNA-seq analyses beyond gene expression. Genome Informatics 5, 10–11.

87. Korsunsky, I., Millard, N., Fan, J., Slowikowski, K., Zhang, F., Wei, K., Baglaenko, Y., Brenner, M., Loh, P.R., and Raychaudhuri, S. (2019). Fast, sensitive and accurate integration of single-cell data with Harmony. Nat Methods 16, 1289–1296. 10.1038/s41592-019-0619-0.

88. Stuart, T., Butler, A., Hoffman, P., Hafemeister, C., Papalexi, E., Mauck, W.M., 3rd, Hao, Y., Stoeckius, M., Smibert, P., and Satija, R. (2019). Comprehensive Integration of Single-Cell Data. Cell 177, 1888–1902 e1821. 10.1016/j.cell.2019.05.031.

89. Thomas, P.D., Ebert, D., Muruganujan, A., Mushayahama, T., Albou, L.P., and Mi, H. (2022). PANTHER: Making genome-scale phylogenetics accessible to all. Protein Sci 31, 8–22. 10.1002/pro.4218.

90. Korotkevich, G., Sukhov, V., Budin, N., Shpak, B., Artyomov, M.N., and Sergushichev, A. (2016). Fast gene set enrichment analysis. BioRxiv, 060012.

91. Robinson, M.D., McCarthy, D.J., and Smyth, G.K. (2010). edgeR: a Bioconductor package for differential expression analysis of digital gene expression data. bioinformatics 26, 139–140.

92. Domcke, S., Hill, A.J., Daza, R.M., Cao, J., O’Day, D.R., Pliner, H.A., Aldinger, K.A., Pokholok, D., Zhang, F., Milbank, J.H., et al. (2020). A human cell atlas of fetal chromatin accessibility. Science 370. 10.1126/science.aba7612.

93. Zhang, Y., Liu, T., Meyer, C.A., Eeckhoute, J., Johnson, D.S., Bernstein, B.E., Nusbaum, C., Myers, R.M., Brown, M., Li, W., and Liu, X.S. (2008). Model-based analysis of ChIP-Seq (MACS). Genome Biol 9, R137. 10.1186/gb-2008-9-9-r137.

94. Bolstad, B.M., and Bolstad, M.B.M. (2013). Package ‘preprocessCore’.

95. Schep, A.N., Wu, B., Buenrostro, J.D., and Greenleaf, W.J. (2017). chromVAR: inferring transcription-factor-associated accessibility from single-cell epigenomic data. Nat Methods 14, 975–978. 10.1038/nmeth.4401.

96. Grant, C.E., Bailey, T.L., and Noble, W.S. (2011). FIMO: scanning for occurrences of a given motif. Bioinformatics 27, 1017–1018. 10.1093/bioinformatics/btr064.

97. Grant, C.E., and Bailey, T.L. (2021). XSTREME: Comprehensive motif analysis of biological sequence datasets. BioRxiv, 2021.2009. 2002.458722.

98. Hao, Y., Stuart, T., Kowalski, M.H., Choudhary, S., Hoffman, P., Hartman, A., Srivastava, A., Molla, G., Madad, S., Fernandez-Granda, C., and Satija, R. (2024). Dictionary learning for integrative, multimodal and scalable single-cell analysis. Nat Biotechnol 42, 293–304. 10.1038/s41587-023-01767-y.

99. Welch, J.D., Kozareva, V., Ferreira, A., Vanderburg, C., Martin, C., and Macosko, E.Z. (2019). Single-Cell Multi-omic Integration Compares and Contrasts Features of Brain Cell Identity. Cell 177, 1873–1887 e1817. 10.1016/j.cell.2019.05.006.

